# Perturbation of the replication stress-ATR axis accelerates human erythroid differentiation but blocks terminal maturation

**DOI:** 10.64898/2026.10.07.754672

**Authors:** Juan Jauregui-Lozano, Amandine Kremer, Karin D. Prummel, Judith B. Zaugg

**Affiliations:** Department of Biomedicine, University of Basel, Basel, Switzerland; Molecular Systems Biology Unit, European Molecular Biology Laboratory, Heidelberg, Germany

**Author notes:** Corresponding authors (J.J-L); (J.B.Z).

## Abstract

Replication stress (RS) can act as an input to differentiation: in erythroid and myeloid cells it is sufficient to accelerate commitment. Yet genotoxic chemotherapy, which imposes replication stress on the bone marrow can cause anemia; a frequent dose-limiting toxicity whose deficit of mature red cells is often attributed to progenitor depletion rather than to any effect on differentiation itself. Here, we ask how RS-induced accelerated commitment gives rise to impaired red cell output, using *in vitro* erythroid differentiation of cord-blood CD34+ progenitors perturbed with clinically relevant chemotherapy agents. We find that treated erythroblasts differentiate faster than controls but fail to mature, measured by lack of enucleated cells. Inhibition of ATR, but not ATM or DNA-PK, reproduces this phenotype. Mechanistically, single-cell profiling of drug-treated cells shows convergence on a transcriptional program, marked by loss of MYB and MYC activity. Treated cultures also accumulate aberrant late erythroblasts depleted for globin and heme programs and enriched for p53 and proteotoxic stress signatures. Together, these results propose that defective maturation is an alternative mechanism for chemotherapy-induced anemia, in addition to depletion of progenitors.

## Introduction

In humans, the bone marrow produces approximately two million erythrocytes every second through an ordered progression from hematopoietic stem and progenitor cells (HSPCs) to committed progenitors, maturing erythroblasts, and enucleated reticulocytes. In contrast to other lineages where cells first exit the cell cycle and subsequently mature, the erythroid precursors do the opposite, acquiring their identity through their terminal divisions. With each successive cell cycle, chromatin condenses progressively, hemoglobin accumulates, and cell volume decreases, culminating in nuclear extrusion (*1*). Erythroid identity is therefore not merely acquired alongside DNA replication but depends on proper cell-cycle progression: G1 shortens progressively through the Colony Forming Unit–Erythroid (CFU-E) stage (*2*), and the transcriptional switch into terminal differentiation is itself S-phase-dependent (*1*, *3*, *4*). Consequently, perturbations to the replication machinery have the potential to alter the differentiation program itself.

A growing body of work has shown that replication stress can act as an input to differentiation in addition to constraining proliferation. In the erythroid and myeloid lineage, nucleotide depletion is sufficient to drive differentiation-associated fate transitions, genotoxic stress biases hematopoietic stem cell behavior in a p53-dependent manner, and relief of a replication-associated block releases leukemic blasts from differentiation arrest (*5*, *6*). Within erythroid cells, folate depletion blocks de novo purine synthesis and drives premature differentiation, proposed as the molecular etiology of folate-deficiency anemia (*7*). Furthermore, hydroxyurea induces fetal hemoglobin through a mechanism separable from its cytoreductive activity (*8*), and erythroleukemic cells activate glycophorin A and hemoglobin expression in response to diverse chemical and metabolic stresses (*9–11*). Altogether, these data establish that a constraint on DNA metabolism can accelerate the erythroid program.

In contrast, clinically relevant genotoxic insults, such as by chemotherapies, are associated to opposing phenotypes: anemia, or lack of mature red blood cells, is amongst the most frequent dose-limiting toxicity of genotoxic therapy (*12*). This is generally attributed to depletion of proliferating progenitors, yet several features of chemotherapy-induced anemia are difficult to reconcile with non-selective killing of cycling cells alone. Anemia predominates in PARP-inhibitor maintenance regimens given without cytoreductive intent (*13*), and is dose-limiting for targeted replication-stress agents such as ATR inhibitors (*14*), in both cases out of proportion to the effect on other proliferative lineages. In mice, a defined block in reticulocyte release accounts for part of 5-fluorouracil-induced anemia (*15*), establishing that an erythroid maturation defect can cause this toxicity. Discriminating progenitor loss from a defect in the differentiation program requires resolution of drug effects along an asynchronous maturation trajectory, which population-level measurements cannot provide.

Here we sought to understand how sublethal genotoxic stress may contribute to aberrant erythropoiesis in a human setting. To do this, we leverage primary cord-blood CD34+ progenitors differentiated *in vitro*, time-resolved flow cytometry and single-cell RNA sequencing and show that mechanistically distinct drugs (hydroxyurea, doxorubicin, topotecan, olaparib) each cause treated cells to progress more rapidly through early maturation but fail to complete enucleation. Inhibition of ATR, but not ATM or DNA-PK, reproduces this phenotype, identifying that the replication-stress-sensing ATR–CHK1 axis, rather than a generalized DNA damage response or progenitor depletion, functions as the basis of this uncoupling of erythroid commitment from terminal maturation. Single cell analysis identified changes to the proliferation networks. Cells that nonetheless reach the orthochromatic stage under treatment occupy a transcriptionally aberrant state, depleted for heme biosynthesis and globin programs and enriched for p53 and proteotoxic stress signatures, rather than completing maturation. Together, these results identify a mechanism by which sublethal replication stress reshapes human erythropoiesis *in vitro*, offering a candidate explanation for forms of chemotherapy-associated anemia that are not accounted for by mere progenitor depletion.

## Results

### 1. A quantitative platform for tracking primary erythroid differentiation through enucleation

To test whether genotoxic chemotherapy perturbs erythroid differentiation under physiologically relevant conditions, we first established a reproducible, biologically faithful readout of erythroid maturation. Specifically, we adapted a well-characterized in vitro CD34+ HSPC differentiation protocol using cord-blood cells in a two-phase culture system, monitored by flow cytometry and scRNA-seq (Fig. 1A). Magnetic enrichment yielded CD34+ purities of 93.9-96.4% across three independent isolations (Supplemental Fig. 1C)

**Figure 1.**
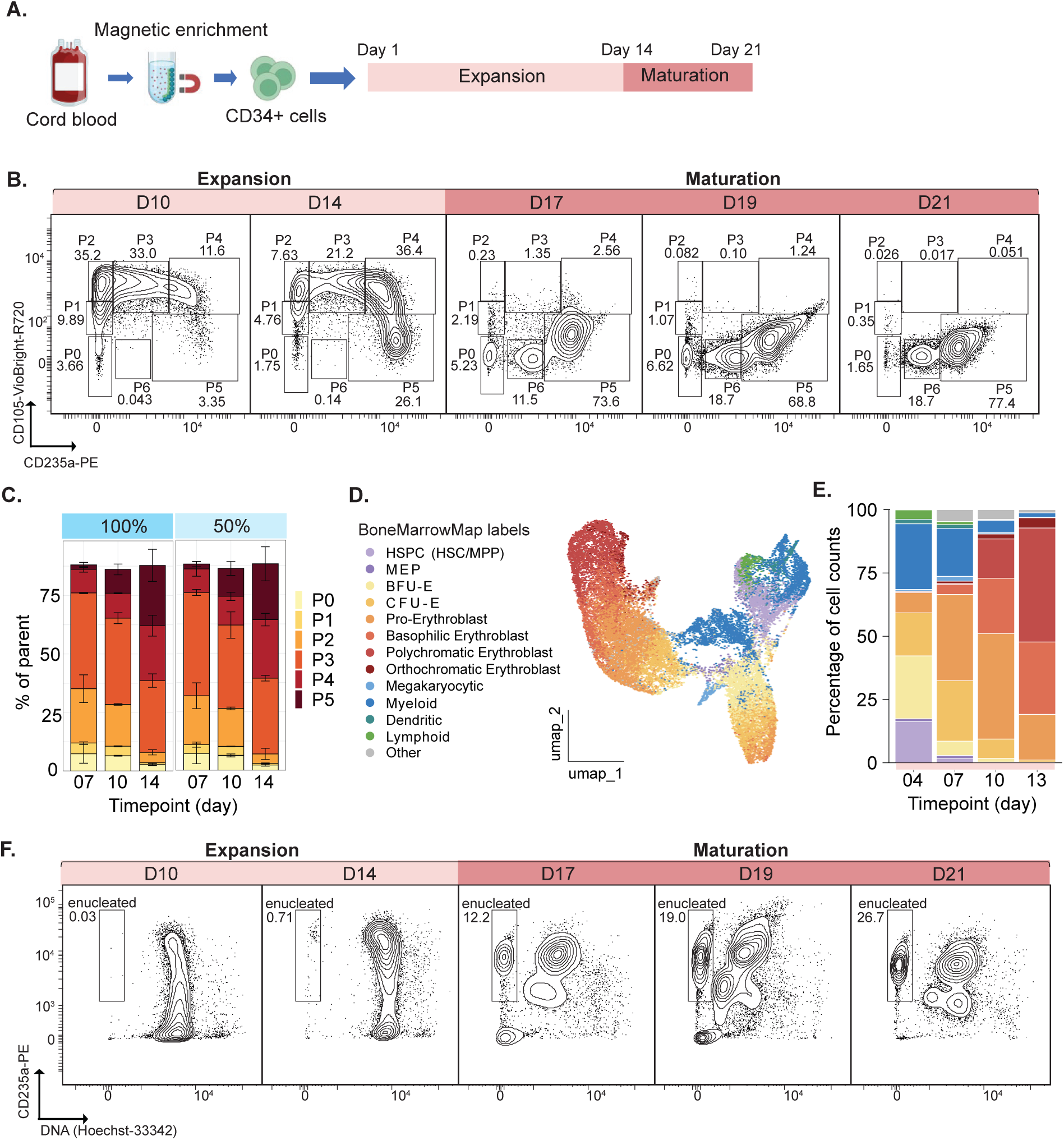
A quantitative platform for tracking primary erythroid differentiation through enucleation. A. Schematic of CD34+ enrichment from cord blood by magnetic selection and the two-phase culture (expansion, day 1–14; maturation, day 14–21) used to generate enucleated cells. B. Representative ENG (CD105-VioBright-R720) versus GYPA (CD235a-PE) contour plots defining sequential maturation gates P0–P6 at days 10, 14 (expansion) and 17, 19, 21 (maturation). Numbers are percentages of the parent gate. C. Quantification of gates P0–P5 at days 7, 10 and 14 in cultures supplemented with full (100%, 1X) or half (50%, 0.5X) erythroid expansion supplement. Bars are mean ± SD of 2 independent cord-blood donors. D. UMAP of the scRNA-seq differentiation time course (days 4–13), colored by BoneMarrowMap-predicted cell type. E. Cell-type composition (percentage of cell counts, BoneMarrowMap labels) at days 4, 7, 10 and 13, colors as in D. F. Representative GYPA (CD235a-PE) versus DNA (Hoechst-33342) plots over the same time course as in B; the gate marks enucleated cells (GYPA+/Hoechst−), with the percentage indicated.

Erythroblast expansion was staged, after exclusion of debris, doublets and dead cells (Supplemental Fig. 1A), using seven sequential flow cytometry gates (P0–P6) defined by endoglin (CD105/ENG) and glycophorin A (CD235a/GYPA) expression (Fig. 1B). These two markers resolve a continuous erythroid maturation trajectory (*16*). Mainly, ENG is transiently expressed across erythropoiesis. It is acquired at the BFU-E to CFU-E transition, peaks in mature CFU-E and proerythroblasts, and is progressively lost from the late basophilic (Baso EB) stage onward. GYPA, by contrast, is acquired at erythroid commitment and maintained throughout. On this basis, the gates correspond to the following stages. P0 (ENG⁻GYPA⁻) contains uncommitted progenitors and comprises all cells at day 1 of culture (Supplemental Figure 1B). P1 (ENG^mid^GYPA⁻) marks erythroid specification, consistent with the BFU-E to immature CFU-E transition. P2 (ENG^high^GYPA⁻) marks peak endoglin, immediately preceding GYPA induction, consistent with mature CFU-E. P3 (ENG^high^GYPA^mid^) corresponds to proerythroblasts committed to terminal differentiation, and P4 (ENG^high^GYPA^high^) to early BasoEBs. P5 (ENG^low^GYPA^high^) spans late basophilic through orthochromatic (OrthoEB) stages. By day 10, 48% of cells are within the GYPA^+^ populations (P3-5). Interestingly, P6 (ENG^⁻^GYPA^low^) emerges only during the maturation phase, present by day 17 of culture, and consistent with post-enucleation pyrenocytes. All quantification below refers to these gates rather than to sorted populations. Gate compositions progressed reproducibly through the expansion phase across two independent cord-blood donors and showed comparable kinetics when using half the standard cytokine supplementation (Fig. 1C), demonstrating that the assay is donor-independent, robust and cost-efficient.

To confirm that the flow-based staging reflects genuine progression through differentiation, we performed scRNA-seq along a differentiation time course of HSPCs during the expansion phase (Days 4–13; quality-control metrics in Supplemental Fig. 1D). Reference mapping to BoneMarrowMap (Supplemental Fig. 1E) and unsupervised clustering resolved the expected erythroid hierarchy with day 4 cells highly enriched for stem progenitor states, such as HSPC and MEP, while erythroblasts maturation stages, such as pro-, basophilic, polychromatic, orthochromatic erythroblasts, dominated at later time points, alongside a minor non-erythroid fraction (megakaryocytic, myeloid, dendritic, lymphoid) (Fig. 1D-E; Supplemental Fig. 1F). The predicted cell-type labels were supported by expression of canonical stage- and lineage-specific marker genes (Supplemental Fig. 1G)

To assess the terminal maturation steps, we quantified enucleated cells (GYPA+/Hoechst-negative) throughout the maturation phase by flow cytometry (Fig. 1F). Importantly, we observed that enucleation rate increased as cells were maintained in maturation media, increasing from 12.2% to 26.7% on day 17 and day 21, respectively. Notably, the enucleated fraction increased progressively, indicating that cells enter enucleation asynchronously across the maturation phase. This makes the fraction of enucleated cells a graded, quantitative readout for assessing drug-induced shifts in terminal maturation.

Collectively, these results establish a quantitative, reproducible platform for tracking primary HSPC erythroid differentiation from early progenitors through enucleated erythroblasts, with enucleation kinetics that recapitulate physiological maturation.

### 2. Mechanistically distinct genotoxic agents accelerate erythroid differentiation and block enucleation at sub-lethal doses

Next we asked whether genotoxic chemotherapy reshapes erythroid differentiation kinetics. We designed a targeted panel of agents that are used clinically and impose replication stress (*17*) through unrelated primary targets: hydroxyurea (ribonucleotide reductase), topotecan (topoisomerase I), doxorubicin (topoisomerase II) and olaparib (PARP1). We then performed a dose-range-finding screen to identify, for each compound, the range of concentrations that reduced cumulative cell expansion over 7 days while leaving viability within 10 percentage points of DMSO and untreated controls (Fig. 2A–B; Supplemental Fig. 2A). We refer to this as the cytostatic window: cells continue to divide and survive, but the culture accumulates fewer cells than control.

**Figure 2.**
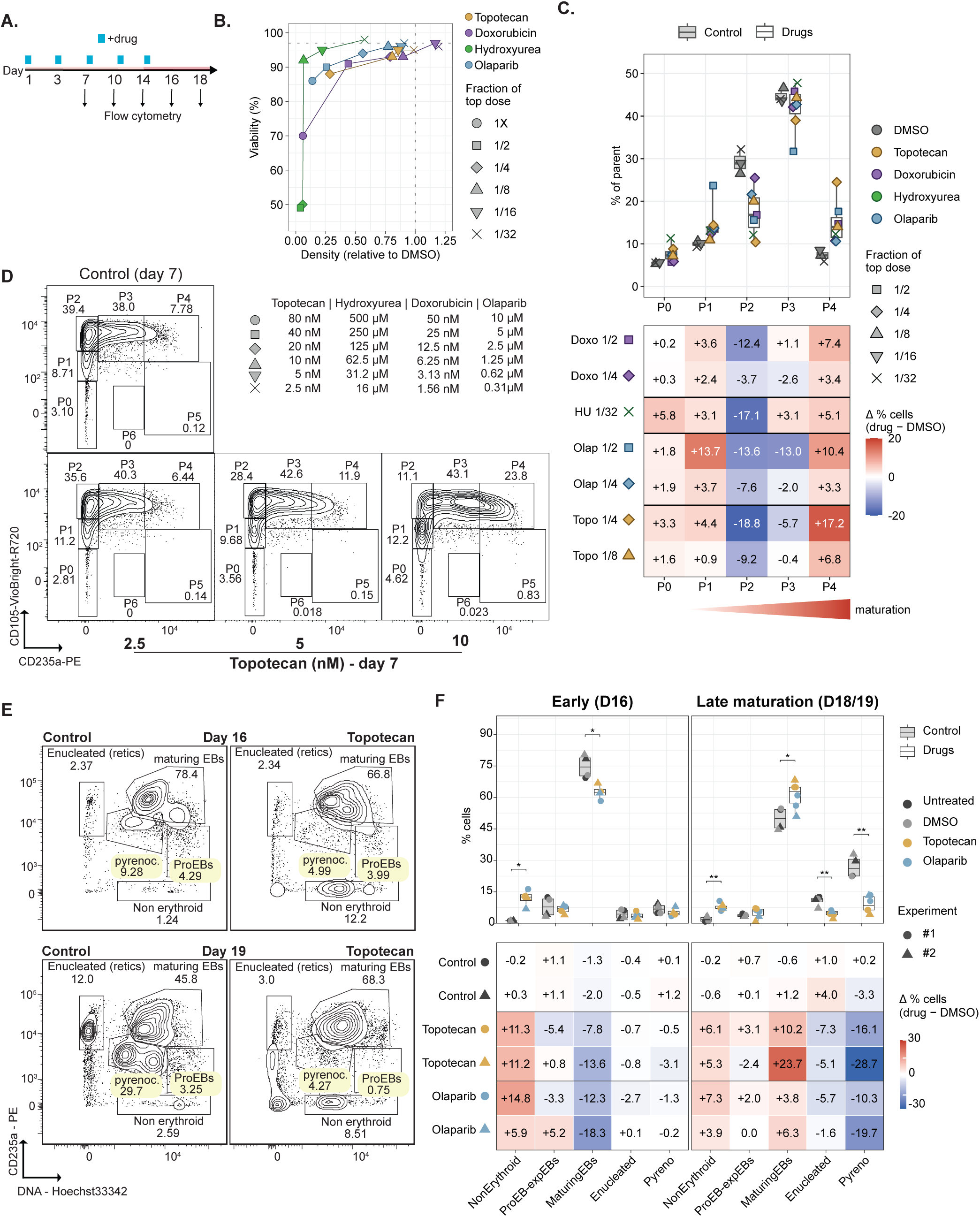
Mechanistically distinct genotoxic agents accelerate erythroid gate transitions and block enucleation. A. Treatment and sampling schedule: drug added from day 1 and refreshed at each medium change (blue marks); flow cytometry at days 7, 10, 14, 16 and 18. B. Culture density relative to DMSO versus viability (% live) at day 7 across a six-point two-fold dilution series (1/32–1X) for topotecan, doxorubicin, hydroxyurea and olaparib. Point size indicates the fraction of the top dose; dashed lines mark the DMSO control. C. Top: distribution of cells across gates P0–P6 at day 7 for controls (DMSO) and the three highest sub-lethal concentrations (#1–#3) of each compound. Bottom: change in the percentage of cells per gate relative to matched DMSO (Δ% = drug − DMSO) for doxorubicin, hydroxyurea, olaparib and topotecan. D. Representative ENG versus GYPA contour plots at day 7 for control and topotecan (2.5, 5, 10 nM); numbers are percentages of parent. E. Representative GYPA versus DNA (Hoechst-33342) gating used in F, control (left) and topotecan (right), at day 16 (top) and day 19 (bottom). Gates define non-erythroid cells, ProEBs, maturing erythroblasts, enucleated cells (reticulocytes) and pyrenocytes. F. Maturation-phase composition at day 16 (early, left) and at a late timepoint, day 18 or 19 (right), across two independent experiments. Top: percentage of cells per population for untreated, DMSO, topotecan and olaparib; symbols denote experiment. Bottom: Δ% versus matched DMSO, one row per compound per independent experiment. Asterisks indicate Wilcoxon test: *p < 0.05, **p < 0.01.

Using these doses, we staged 7-day cultures by ENG and GYPA (Fig. 2C). Every compound moved cells out of the CFU-E compartment and into GYPA-expressing stages earlier than controls, in a dose-dependent manner: relative to DMSO, the proportion of cells in P2 decreased by 9.2 percentage points at 10 nM topotecan and by 18.8 at 20 nM, with comparable shifts for olaparib, hydroxyurea and doxorubicin (Fig. 2-C; Supplemental Fig. 2B). The cells leaving the P2 fraction were not lost from the culture but redistributed forward: every compound that reduced the P2 compartment increased the GYPA-high P4 fraction (+3.3 to +17.2 percentage points), and the combined GYPA+ fraction increased in six of the seven conditions. Representative gating illustrates this redistribution directly across a narrow range of topotecan (Fig. 2D). Because GYPA is acquired at erythroid commitment and retained thereafter, a larger GYPA+ compartment at a fixed timepoint means a greater proportion of cells had progressed beyond erythroid commitment by day 7 relative to controls.

We next asked whether this accelerated early commitment translates into more efficient terminal maturation using GYPA and DNA staining(see figure 1F). Terminal maturation experiments require extended culture and are lower throughput, so we focused on two agents chosen to represent complementary routes to replication stress: topotecan, which generates lesions directly by trapping topoisomerase I cleavage complexes ahead of the replisome, and olaparib, which does not itself damage DNA but traps PARP1 on endogenous single-strand breaks and prevents their resolution. We followed drug treated cultures sampling during early and late maturation. Strikingly, we found that at day 16, drug-treated cells showed a depletion of maturing erythroblasts, with a concomitant accumulation of non-erythroid cells (as determined by alive GYPA-DNA+ cells) (Figure 2E-F). This is consistent with a reduced erythroid expansion that fails to dilute the non-erythroid compartment. By the later timepoint, the EB population decreased again in control cells, giving rise to pyrenocytes and enucleated cells, while the drug-treated cells failed to mature, as shown by an aberrant accumulation of maturing EBs (Figure 2E-F). Because a single enucleation event yields both an enucleated cell and its pyrenocyte, this coordinated loss of both downstream populations alongside upstream maturing EB accumulation indicates that drug-treated erythroblasts fail to enucleate.

Together, these data describe a two-part phenotype in which sublethal genotoxic stress, regardless of the insult or molecular target, first accelerates erythroid cells through early differentiation at the expense of expansion, and then causes cells to stall before completing enucleation.

### 3. Loss of ATR, but not ATM or DNA-PK, is sufficient to induce accelerated and impaired erythroid differentiation

Given the phenotypic convergence of mechanistically distinct genotoxic drugs, we next asked which arm of the damage response translates genotoxic stress into an erythroid differentiation phenotype. In myeloid models, inhibiting checkpoint signaling does not prevent replication stress-induced differentiation, indicating that checkpoint signaling is not the transducer of the differentiation signal (*5*). In the same study, however, ATR inhibition on its own was sufficient to induce myeloid markers. Together, these observations are consistent with the checkpoint acting as a buffer that keeps replication stress below the level at which it alters cell fate. We therefore reasoned that if replication stress exceeding checkpoint capacity is the shared driver, lowering checkpoint capacity should reproduce the drug-induced phenotype without exogenous damage, and that this should be specific to the replication-stress-sensing arm.

Genotoxic lesions are sensed and transduced by three related PIKK-family kinases: ATR, ATM and DNA-PK. ATR is activated by single-stranded DNA at stalled or collapsed replication forks and signals through CHK1 (*17*, *18*). ATM and DNA-PK are both recruited to double-strand breaks, but differ in output: ATM signals a checkpoint through CHK2, whereas DNA-PK drives non-homologous end joining (*17*) (Fig. 3A). We therefore treated differentiating CD34+ cultures with selective inhibitors of each kinase and identified the cytostatic range at which cells decreased proliferation while viability remained within 10 points of control, at day 7 of cultures (Fig. 3B; Supplemental Fig. 3A). Notably, we found that both ATRi (Ceralasertib) and DNA-PKi (Nedisertib) treatment showed a dose-dependent response in viability and density, while ATMi treatment (AZD1390) led to dose-dependent loss of proliferation but kept cells viable regardless of the concentration (Fig. 3B), suggesting that expansion of erythroid progenitors tolerate loss of ATM signaling but not loss of replication-stress buffering or end-joining capacity.

**Figure 3.**
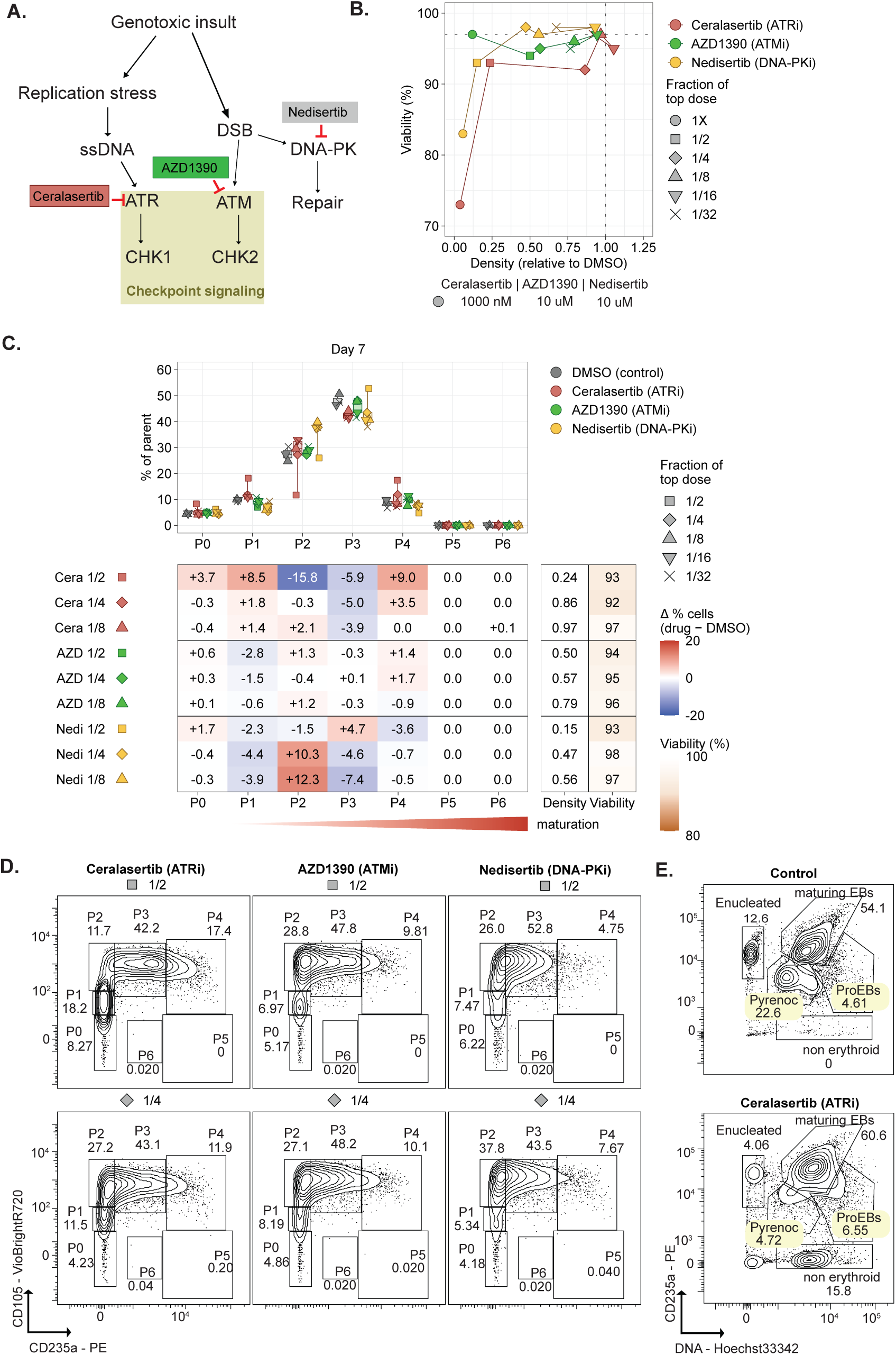
Loss of ATR, but not ATM or DNA-PK, is sufficient to accelerate erythroid gate transitions and impair maturation. A. Schematic of checkpoint signalling downstream of genotoxic insult: replication stress and ssDNA signal through ATR–CHK1 (inhibited by ceralasertib), double-strand breaks (DSB) through ATM–CHK2 (AZD1390) and DNA-PK-dependent repair (nedisertib). B. Viability (% live) versus culture density relative to DMSO at day 7 across a six-point two-fold dilution series (1/32–1X) of ceralasertib (ATRi), AZD1390 (ATMi) and nedisertib (DNA-PKi). Dashed lines mark the DMSO control. C. Top: distribution of cells across gates P0–P6 at day 7 for each inhibitor (colored) versus DMSO (grey); symbols denote the fraction of the top dose. Bottom: change in the percentage of cells per gate relative to matched DMSO (Δ% = drug − DMSO) at each dose (0.5X–0.032X). D. Representative ENG versus GYPA contour plots at day 7 for ceralasertib, AZD1390 and nedisertib at two doses each. Numbers are percentages of parent. Compare with the DMSO control in Fig. 2D. E. Representative GYPA versus DNA (Hoechst-33342) plots at the late maturation timepoint (day 18) for control (top) and ceralasertib (bottom), gated as in Fig. 2E.

Next, we analyzed differentiation kinetics of the cultures. ATR inhibition reproduced the accelerated differentiation associated with genotoxic drugs, shifting cells out of the “mature CFU-E” population P2 and into GYPA-expressing stages (P2 −15.8 percentage points, P4 +9.0 at the highest dose). ATM inhibition produced no comparable shift at any concentration tested, despite restraining proliferation. DNA-PK inhibition moved cells in the opposite direction, increasing late CFU-E enriched P2 by 10 to 12 percentage points across most of the range (Fig. 3C-D). Altogether, these data indicate that accelerated erythroid differentiation tracks specifically with loss of replication-stress signaling rather than with damage signaling in general or with restrained proliferation. Neither double-strand-break-directed perturbation (ATMi, DNA-PKi) reproduced the phenotype. Furthermore ATMi restrained proliferation to a comparable degree without moving cells forward, arguing that slowed expansion alone is not sufficient to shift erythroid kinetics.

We next asked whether ATRi was sufficient to impair maturation of erythroblasts. By the late maturation timepoint, ATRi reproduced the maturation defect as well. Maturing erythroblasts accumulated (+5.9 and +9.4 percentage points in two independent experiments while both enucleated cells and pyrenocytes decreased (-7.5 and -2.5, and -17.7 and -23.9, respectively; Fig. 3E; Supplemental Fig. 3B), the same coordinated loss of both enucleation products observed with topotecan and olaparib (see Fig. 2E-F). Inhibition of ATR alone, without exogenous DNA damage, is therefore sufficient to reproduce the accelerated yet impaired differentiation observed in erythroblasts undergoing differentiation in the presence of genotoxic drugs.

### 4. Three agents with distinct targets converge on a near-identical transcriptional response

We next sought to dissect how perturbation of the ATR axis remodels erythroid differentiation transcriptionally. Because treated and control cultures differ in stage composition, transcriptional analysis requires comparing matched cell types rather than bulk populations. We therefore profiled cultures treated with topotecan, olaparib and ceralasertib, two agents imposing replication stress upstream of ATR and one inhibiting it directly, by scRNA-seq, across three expansion timepoints and one maturation timepoint (Supplemental Fig. 4A). After initial QC (Supplemental Fig. 4B), we obtained 82,275 high quality cells (cells captured per sample in Supplemental Fig. 4C). To obtain cell type annotations, we mapped the cells to the BoneMarrowMap reference (*19*), which yielded the expected distribution of cell types along the erythroid trajectory (Fig. 4A), with accumulation of mature cell types with the later time points (Supplemental Fig. 4D). Notably, all drugs pushed the cells along a distinct trajectory that diverged from the control cells between CFU-E and Basophilic Erythroblasts (Baso EB) stage and converged back at the Polychromatic Erythroblast (Poly EB) stage (Fig. 4B; Supplemental Fig. 4E). Unsupervised clustering analysis identified 15 clusters (0–14), of which none were specific to one drug (Fig. 4C). Rather, we found clusters with either an equal representation of the five conditions (such as Cluster 2, 3, 5, and 7), with mostly control cells (Cluster 1 and 4) or clusters that had equal representation from all three drug treatments (Cluster 0, and 11). The changes in cell type composition and stage-specific differential expression relative to controls were almost identical across drugs (Supplemental Fig. 4F, Fig. 4D)(olaparib and topotecan R² = 0.91, 97% concordant sign; ceralasertib and olaparib R² = 0.66, 91%; ceralasertib and topotecan R² = 0.67, 92%; Fig. 4D - top), despite extensive and stage-dependent differential gene expression between each of the drugs and controls, peaking at the pro-erythroblast stage (3,477–5,319 Differentially Expressed Genes (DEGs) per drug) and falling sharply at the orthochromatic stage (177–997; Fig. 4E). Fold changes were consistent regardless of comparison (untreated or DMSO-matched, Fig 4D - bottom). Altogether, these analyses demonstrate that sustained replication stress redistributes erythroblasts undergoing differentiation into a transcriptionally distinct profile compared to controls.

**Figure 4.**
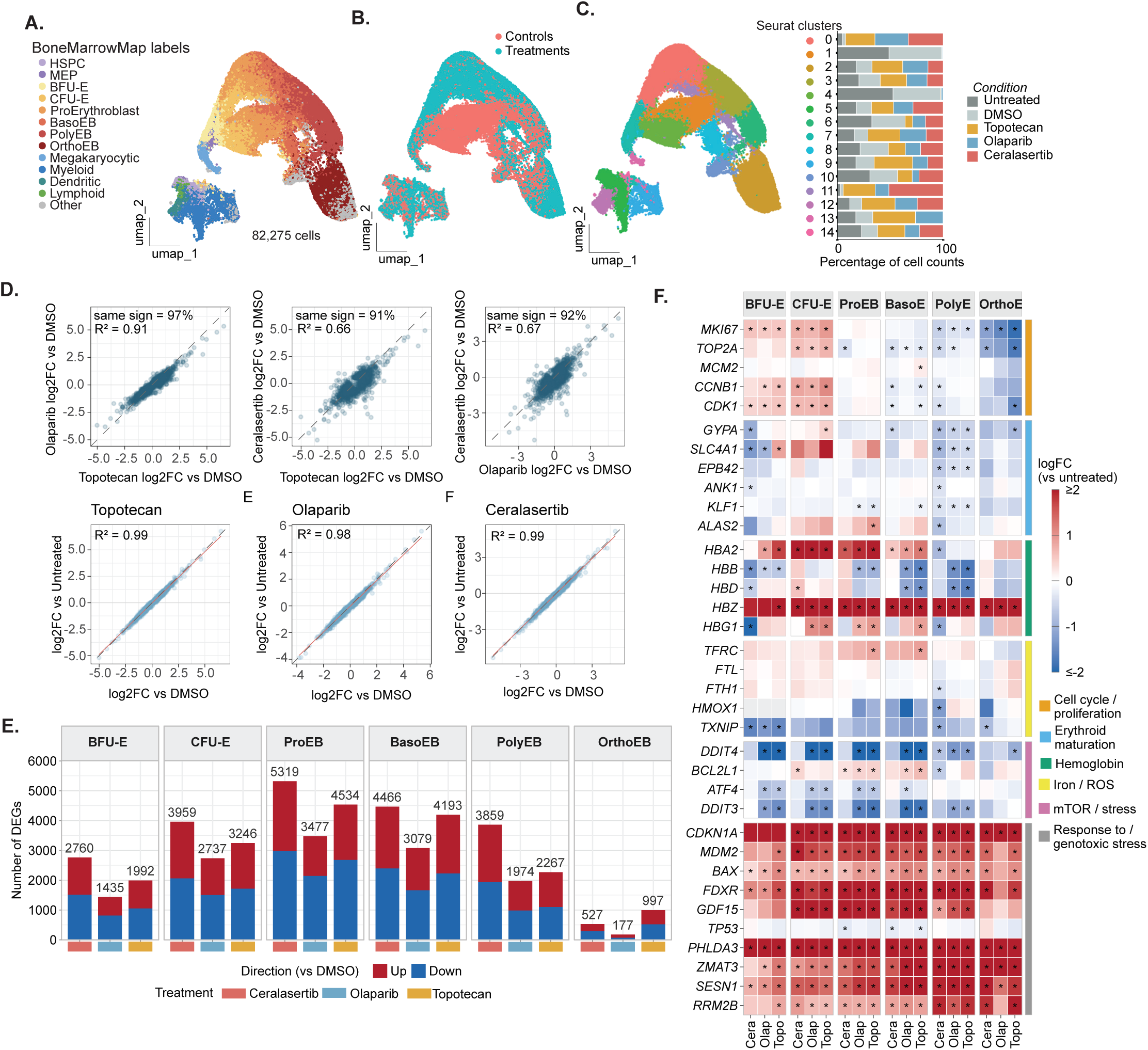
Three agents with distinct targets converge on a near-identical transcriptional response. A. UMAP of 82,275 cells from cord-blood CD34+ HSPCs differentiated under five conditions (untreated, DMSO, topotecan, olaparib, ceralasertib) and sampled at days 7, 10, 13 and 16, colored by BoneMarrowMap-predicted cell type. B. The same embedding colored by group (Controls = untreated + DMSO; Treatments = topotecan + olaparib + ceralasertib). C. Left: the same embedding colored by Seurat cluster (0–14). Right: condition composition of each cluster as a percentage of cell counts. D. Pairwise comparison of stage-resolved log2 fold changes (log2FC). Top: between drugs (olaparib versus topotecan; ceralasertib versus topotecan; ceralasertib versus olaparib), with R² and the percentage of genes changing in the same direction. Bottom: for each drug, log2FC versus untreated plotted against log2FC versus DMSO, showing insensitivity to the choice of control. E. Number of differentially expressed genes per erythroid stage and treatment (versus DMSO), split by direction of change. F. Stage-resolved log2 fold change (versus untreated) for curated gene sets covering cell cycle/proliferation, erythroid maturation, hemoglobin, iron/ROS, mTOR/stress and the response to genotoxic stress. Asterisks mark significance (adj. p < 0.05) established by Seurat’s *findAllMarkers* function: Wilcoxon Rank Sum test.

Further inspection of DEGs (Fig. 4F) revealed a biphasic drug-response of the proliferation genes (*MKI67, TOP2A, MCM2*), with an initial upregulation during BFU-E and CFU-E stages, and a latter downregulation across more committed erythroid stages (BasoEB – OrthoEB), suggesting dynamic dysregulation of cell cycle gene programs throughout differentiation, co-occurring with sustained reduced proliferation (see Fig 2 and 3). Canonical erythroid maturation genes (*GYPA, SLC4A1, EPB42, ANK1, KLF1, ALAS2*) showed mixed but predominantly negative fold changes at later stages (PolyEB, OrthoEB), indicating lack of induction of the terminal maturation machinery. Hemoglobin genes (*HBA2, HBB, HBD, HBZ*) showed stage-dependent modulation, with the largest changes relative to controls concentrated at Pro-Erythroblast through PolyEB. Genes that normally respond to genotoxic stress, such as *CDKN1A, MDM2, BAX, FDXR and GDF15,* showed a strong consistent upregulation across nearly all stages and all three drugs – with the notable exception of TP53, which showed no change in expression.

Together these data show that three agents with unrelated primary targets converge on a single transcriptional response, concentrated in the progenitor compartment, in which cell-cycle genes are dysregulated in a stage-dependent manner, there is a consistent stress response, and the terminal maturation machinery is induced in different cell types relative to controls.

### 5. Treatment collapses the MYB–MYC proliferative network and imposes a repressive-E2F state

As cell-cycle genes were among the most consistently affected across stages, we next examined proliferation, starting with culture density and viability over the full time course. Analysis of density and viability of the cultures throughout expansion and maturation revealed that, relative to controls, drug-treated cells expanded less from day 7 onward, with topotecan- and ceralasertib-treated cultures reaching roughly two logs less cumulative expansion than control by day 16 (Fig. 5A, top) and viability collapsing in the maturation phase, two days after the switch into maturation medium (Fig. 5A, bottom). Further, the expansion deficit was accompanied by a shift in cell-cycle occupancy rather than a uniform slowing, as measured by cell cycle phase scoring using canonical cell cycle transcriptional markers (*20*) (Fig. 5B). Drug-treated cells showed a depletion of G1 in BFU-E and CFU-E (-14.3 and -6.9 percentage points, respectively) with a corresponding increase of G2/M at these stages (+8.5 and +9.8 points), indicating a partial G2/M accumulation at the progenitor stages rather than a simple G1 shift. For later stages, this reversed: BasoEB populations showed a G1 increase (+15.6 points) and G2M/S-phase depletion (-5.7 and -9.9 points), suggesting that cells reaching these stages later in the treatment course stall in G1 rather than progress through S phase. Later stages (PolyEB, OrthoEB) showed minimal cell-cycle phase redistribution. Phase distributions for each of the five conditions separately are shown in Supplemental Fig. 5A; untreated and DMSO cultures were near-identical at every stage).

**Figure 5.**
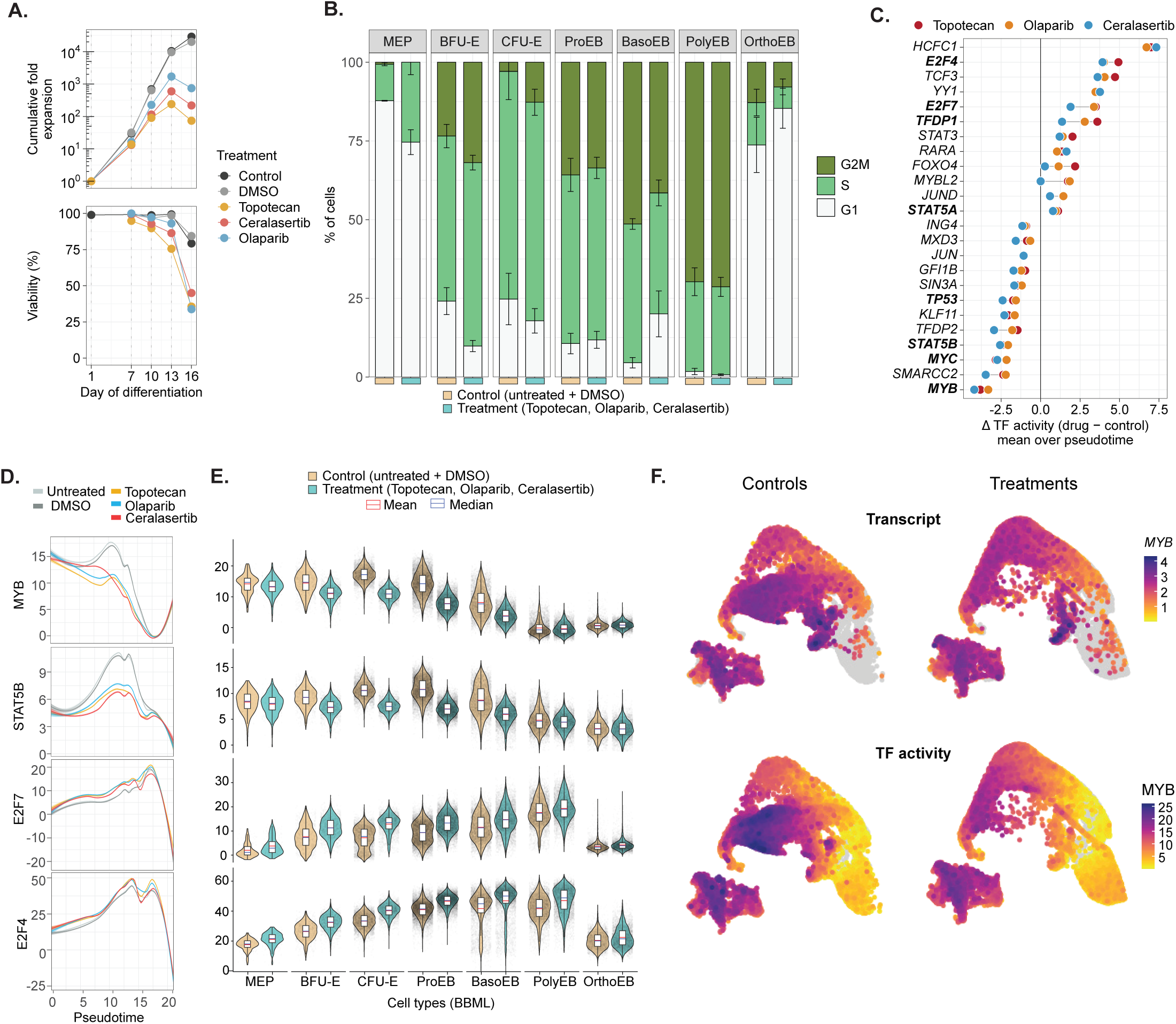
Treatment collapses the MYB–MYC proliferative network and imposes a repressive-E2F state. A. Cumulative fold expansion (top) and viability (bottom) across the differentiation time course (days 1–16) for each condition. B. Percentage of cells in G1, S and G2/M per BoneMarrowMap cell type, control (untreated + DMSO) versus treatment (topotecan, olaparib, ceralasertib). Bars are mean ± SD across timepoints. C. Difference in mean per-cell regulon activity between drug and control, averaged over erythroid pseudotime, for the 24 most differentially active transcription factors; one point per drug. D. MYB, STAT5B, E2F7 and E2F4 regulon activity along pseudotime, by condition. E. Per-cell-type distributions of the same four regulons, control (untreated + DMSO) versus treatment; red and blue marks indicate mean and median. F. MYB transcript (top) and MYB regulon activity (bottom) projected on the UMAP, controls versus treatments.

Neither the expansion deficit nor the stage-dependent phase redistribution was matched by downregulation of cell cycle-associated genes; they were induced at the BFU-E and CFU-E stages (see Fig. 4F). To identify the transcriptional regulators underlying these changes in cell-cycle gene expression, we quantified differential transcription factor (TF) activity between drugs and control. For this, we inferred gene regulatory networks (GRNs) with pySCENIC (*21*), which assign each TF a regulon: the set of target genes that are co-expressed with it and carry its binding motif (773 regulons; Supplemental Fig. 5B-C). We then calculated per-cell TF activity based on the expression of the genes in their regulon, using decoupleR (*22*), and refer to this as regulon activity. For each TF, we binned cells along erythroid pseudotime, computed the difference in mean regulon activity between each drug and control within each bin, and averaged these differences across bins. This identified several drug-responsive TFs. Across all three drugs the largest decrease in activity was in erythroid-enriched MYB, followed by SMARCC2, MYC, STAT5B, TFDP2 and KLF11, with smaller reductions in GFI1B, SIN3A, JUN and MXD3 (Fig. 5C). On the opposite end, HCFC1, E2F4, TCF3, YY1, E2F7 and TFDP1 showed increased regulon activity. The three drugs agreed on both the identity and the rank order of these changes, extending the phenotypic convergence to the level of regulatory networks.

MYB activity was reduced across the whole trajectory, with the gap widening at the MEP-to-CFU-E transition and persisting through the basophilic stage before narrowing as cells approach terminal maturation (Fig. 5D, E). Projecting MYB expression and MYB regulon activity separately onto the UMAP showed that the loss of activity is broader than the loss of transcript, indicating a change in regulon output rather than simple downregulation of the factor (Fig. 5F). Interestingly, MYB controls amplifying divisions of erythroid progenitors, and its partial knockdown in human CD34+ cells produces precocious erythroid differentiation, slowed cell-cycle progression and increased representation of non-erythroid lineages (*23*), the same characteristics we report by flow cytometry and by composition analysis. The concurrent loss of STAT5B activity points the same way: loss of STAT5 signaling in erythroblasts drives differentiation at the expense of renewal, with premature proliferation arrest (*24*).

All three drugs also increased the activity of the regulons assigned to E2F4 and E2F7 (Fig. 5C–E). Both factors are transcriptional repressors: E2F4 silences cell-cycle genes during cell-cycle exit, and E2F7 is an atypical repressor required for the 4N-G1 arrest that follows genotoxic stress (*25*). Because regulon activity is calculated from the expression of target genes, higher activity of a repressor’s regulon means that its targets are more highly expressed, indicating reduced repression. Consistent with this, canonical E2F targets were induced in the progenitor compartment (Supplemental Fig. 5D). This is in line with a model in which the replication-stress checkpoint sustains cell-cycle transcription: CHK1, activated downstream of ATR, phosphorylates E2F6 (*26*) and E2F7/8 (*27*), and thereby inhibits their repressor function. Topotecan and olaparib, which activate this checkpoint, would be expected to derepress E2F targets by this route. Ceralasertib produced the same change although ATR inhibition should instead release these repressors, so a single mechanism common to all three drugs cannot be assigned from these data. However, reduced MYB activity and derepressed E2F targets in progenitors are compatible with a shortened phase of amplifying divisions, which may contribute to the maturation stall measured by flow cytometry.

### 6. Treated cells accumulate in an aberrant late erythroblast state that is reproducible across independent datasets

Having established that drug treatment induces a transcriptional program marked by elevated stress signaling and dysregulated proliferation, we next asked whether these changes could account for the downstream defect in enucleation observed by flow cytometry. Composition analysis at the later time points revealed a progressive increase in cells that BoneMarrowMap could not confidently assign to any reference state, labelled "Other", which co-occurred with a depletion of the late erythroid stages, mainly polychromatic and orthochromatic erythroblasts, in addition to accumulation of non-erythroid cells (Fig. 6A and Supplemental Fig. 4F), reaching up to 50% of topotecan-treated cells by Day 16. Inspection of cell type distribution showed that these “Other” cells were accumulating in Cluster 2, which was OrthoEB specific, and that Cluster 2-associated “Other” cells were heavily enriched in drug-treated cells, unlike the “Others” cells in Cluster 9 and 10. Re-embedding of the OrthoEB-enriched Cluster 2 cells separated cells by condition (drug, control) and annotation status (passed annotation – PASS, failed annotation to BoneMarrowMap – FAIL) (Fig. 6B-C).

**Figure 6.**
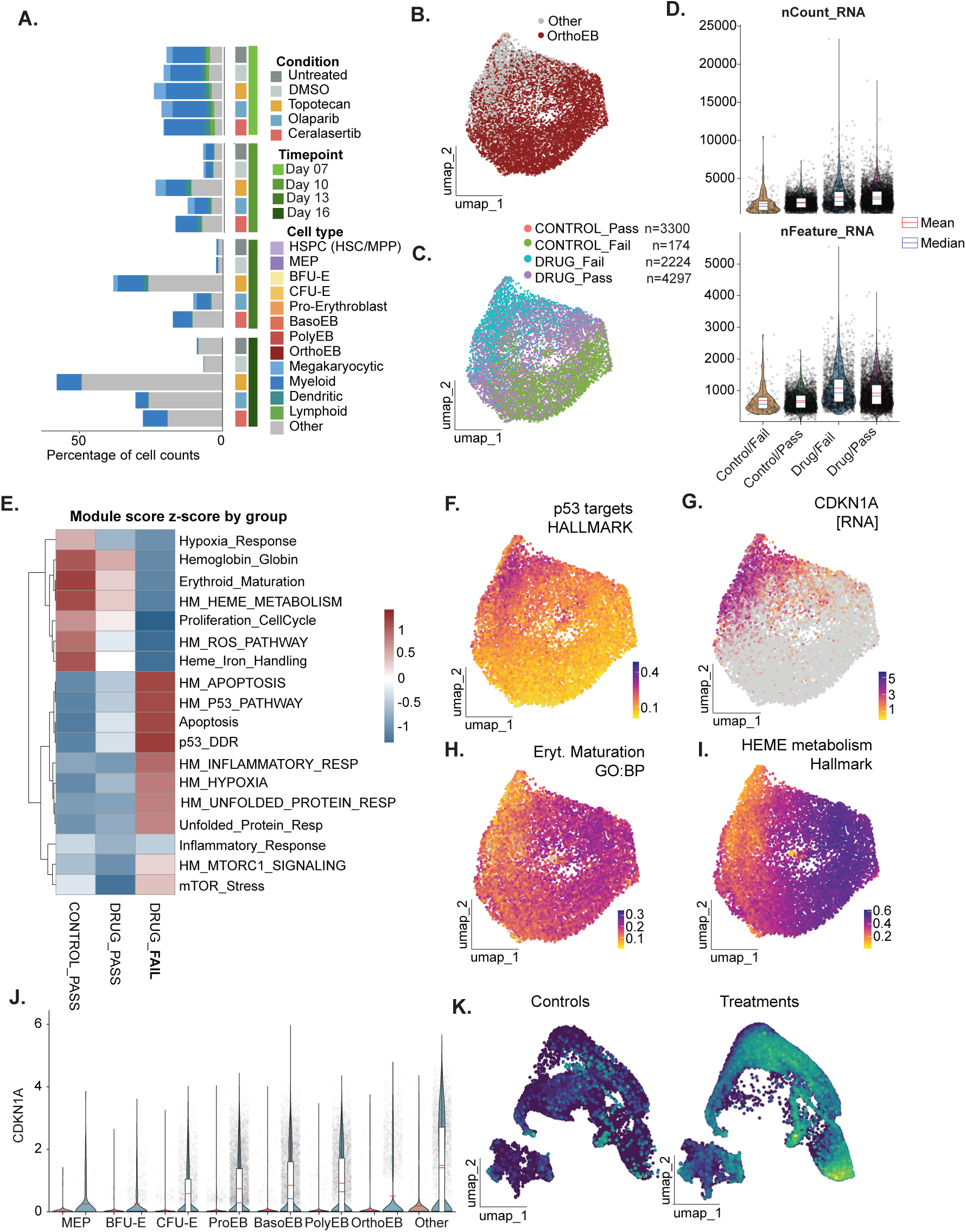
Treated cells accumulate in an aberrant late erythroblast state. A. Cell-type composition (percentage of cell counts, BoneMarrowMap labels) by condition and timepoint; the grey "Other" fraction increases in drug-treated cultures at later timepoints. B. UMAP of the orthochromatic-erythroblast-associated cluster 2 after re-embedding, colored by BoneMarrowMap label (OrthoEB versus "Other"). C. The same embedding colored by the four combinations of group and BoneMarrowMap mapping quality (CONTROL_Pass n = 3,300; CONTROL_Fail n = 174; DRUG_Fail n = 2,224; DRUG_Pass n = 4,297); Fail cells occupy a contiguous region rather than scattering. D. nCount_RNA (top) and nFeature_RNA (bottom) per group; red and blue marks indicate mean and median. DRUG_Fail cells show the highest values, arguing against a low-complexity technical artefact. E. Mean module-score z-scores across CONTROL_PASS, DRUG_PASS and DRUG_FAIL for MSigDB hallmark and curated erythroid/stress gene sets, hierarchically clustered by row. F–I. Module scores and expression projected on the cluster-2 re-embedding: p53 targets hallmark module (F), CDKN1A expression (G), erythroid maturation (GO:BP) module (H) and heme metabolism hallmark module (I). J. CDKN1A expression by cell type, control versus treatment; red and blue marks indicate mean and median. K. CDKN1A expression projected on the full-dataset UMAP, controls versus treatments.

To exclude that the FAIL cells simply represent low-quality, low-complexity cells we compared the counts (nCount_RNA) and (nFeature_RNA) per cell. FAIL cells from the drug-treated experiments showed the highest nCount_RNA and nFeature_RNA among all four groups, thus arguing against low-quality cells (Fig. 6D), and instead supporting a distinct transcriptional state induced by genotoxic drugs.

To characterize the drug-specific FAIL population transcriptionally, we scored cells for a panel of curated and HALLMARK gene modules and compared mean z-scores across control-PASS, drug-PASS, and drug-FAIL groups (Fig. 6E). This revealed that erythroid-identity modules were most enriched in control-PASS and progressively reduced in drug-treated cells with drug-FAIL cells showing the strongest depletion. Conversely, stress- and death-associated modules were low in control-PASS but markedly elevated specifically in drug-FAIL, with drug-PASS generally intermediate or unchanged relative to control. This pattern indicates that drug-treated cells that didn’t map to any known bone marrow population adopt a transcriptional state combining loss of erythroid maturation and proliferative programs with activation of p53-driven apoptotic and proteotoxic/inflammatory stress responses.

Altogether, these results indicate that sustained replication stress reshapes terminal erythropoiesis by generating an alternate late-erythroblast state, characterized by attenuated erythroid maturation programs and persistent stress signaling, rather than by simply blocking or delaying normal maturation. Notably, both drug-PASS and drug-FAIL cells co-occupy the same Seurat cluster (Cluster 2), indicating that despite their divergent mapping outcomes, these cells remain transcriptionally more similar to one another than to any other population in the dataset.

Projecting module scores of the selected gene lists onto the re-embedded Cluster 2 UMAP revealed that they cluster separately: We found that there was enrichment for the TP53 hallmark genes (Fig. 6F), almost exclusively enriched amongst the cells that failed to map to the reference. For example, p53 effector p21 (or CDKN1A) was exclusively enriched in the FAIL cells (Fig. 6G). Further, Erythroid maturation and Heme metabolism associated scores were high across most of the cluster but sharply dropped off in a discrete subregion of the cluster where drug-FAIL cells accumulate (Fig. 6H-I). This data indicates that loss of erythroid maturation and activation of p53-related stress responses co-localize to these FAIL cells. Having observed consistent *CDKN1A* upregulation across cell types in drug-treated cultures (See Fig. 4F), we next examined CDKN1A expression across the full erythroid differentiation trajectory in control and drug treated cells (Fig. 6J-K). Strikingly, we found that control cells upregulate CDKN1A mildly in middle stage progenitors but show no expression at all in the later maturation stages. However, drug-treated cells sustained CDKN1A through terminal stages and this persistence was specific to the FAIL cells (Fig. 6J-K). In contrast, drug-treated cells that mapped to the OrthoEB suppressed CDKN1A to control levels. This persistent CDKN1A expression points to a sustained targeted checkpoint arrest rather than a generalized p53 transcriptional response, and may represent the terminal transcriptional signature of cells unable to resolve replication stress accumulated earlier in differentiation.

Finally, we asked whether the off-reference population arises independently of our specific protocol. We applied the same PASS/FAIL classification to a published dataset in which adult CD34+ HSPCs were differentiated towards the erythroid lineage under the replication-stress agents hydroxyurea (HU) and aphidicolin (APH) (*5*). Drug-treated cells showed a progressive, timepoint-dependent accumulation of “Other”-classified cells, mirroring the pattern we observed in our own dataset (Supplemental Fig. 6A). As in our dataset, these cells did not show reduced library complexity relative to assigned cells (Supplemental Fig. 6B), and drug-treated FAIL cells combined depletion of erythroid-maturation and globin modules with elevated p53 and stress-response signatures (Supplemental Fig. 6C). Notably, this accumulation was already evident by Day 14 in the HU/APH dataset, the latest timepoint captured by their differentiation protocol, which was not carried through to terminal maturation and enucleation. Regardless, this suggests that the emergence of the FAIL population begins during earlier stages of terminal erythroid differentiation and is not merely a consequence of prolonged culture in our own maturation-extended protocol.

## Discussion

We set out to test whether cytostatic concentrations of mechanistically diverse genotoxic drugs converge on a common effect on erythropoiesis, and to identify the pathway responsible. In primary CD34+ hematopoietic stem and progenitor cells, four drugs with distinct primary targets, ribonucleotide reductase, topoisomerases I and II, and PARP, produced a single convergent phenotype: accelerated erythroid commitment and differentiation coupled to a failure of terminal maturation and enucleation. This phenotype was reproduced by inhibition of ATR but not by inhibition of ATM, even at doses that restrained proliferation more completely and with no loss of viability, identifying the replication-stress-sensing ATR-CHK1 axis and the loss of checkpoint restraint rather than growth arrest itself as its central driver. At the level of regulatory state the change that tracks with the phenotype is a repression of MYB and MYC transcriptional activity with reciprocal gain of repressive E2F activity in the progenitor compartment (by reducing its repressive function), rather than precocious activation of the erythroid master regulators. Together these results argue that a substantial component of chemotherapy-associated anemia is not simply the loss of proliferating progenitors, but an active reprogramming of the erythroid differentiation trajectory by replication stress.

The most striking feature of the phenotype is that the same perturbation speeds early differentiation yet blocks its completion. We interpret this as an uncoupling of two normally linked processes: the transcriptional commitment to the erythroid program and the terminal maturation divisions that must complete before enucleation. Erythroid identity is not acquired before the terminal divisions but through them, and the switch into terminal differentiation is itself S-phase-dependent(*1*), so a perturbation that shortens or disrupts the amplifying phase advances commitment and depletes division capacity. MYB sets the number of amplifying divisions a progenitor completes, and MYB-low states produce accelerated but impaired differentiation with slowed cycling and lineage output (*23*). Our data is thus consistent with a model in which maturation advances ahead of division history resulting in fewer divisions of the treated cells when reaching the orthochromatic compartment.

All the drugs tested in our study share the capacity to provoke replication stress, through nucleotide depletion, trapped topoisomerase complexes, direct ATR inhibition, or PARP trapping and unrepaired replication gaps. The ATR-versus-ATM dissection confirms that it is the sensing of single-stranded DNA and stalled forks, not of double-strand breaks, that reshapes the erythroid trajectory. ATM is largely dispensable for steady-state erythropoiesis in young animals; the hematopoietic failure that eventually develops in Atm⁻/⁻ mice reflects loss of HSC self-renewal under oxidative stress rather than a defect in erythroblast maturation (*28*). The primary surveillance pathway in erythroblasts seems to be ATR-CHK1: erythroblasts depend on PARP2 activity to ligate the Okazaki fragment nicks that accumulate during the ultra-fast replication forks of terminal erythroid divisions (*29*), and loss of this tolerance engages CHK1 and RPA phosphorylation as a first-line response (*30*), with p53-dependent apoptosis following only when this mechanism fails.

A plausible mechanistic bridge between the accelerated commitment and the maturation stall comes from a recent study of ATR inhibition in non-malignant epithelial cells, which identified MDM2 and CDKN1A as the top nascent-transcription targets induced downstream of ATR pathway loss and showed that this p53-p21 output drives a p53-p21-RB1-dependent exit from the cell cycle in G2 rather than completion of mitosis (*31*). Because terminal erythroid maturation requires a defined series of terminal divisions prior to enucleation, an analogous RB1-dependent G2 exit in basophilic and polychromatic erythroblasts offers a candidate explanation for the stall we observe. Our regulon data support this bridge from the transcriptional side: E2F4 is the principal partner of RB-family pocket proteins in repressive complexes that silence cell-cycle genes during exit, and E2F7 is required for cells to execute a 4N-G1 arrest after genotoxic stress (*25*).

Clinically, this pathway-defined mechanism has two practical consequences. First, erythroid output can be compromised even when progenitor numbers are relatively preserved, so standard measures of marrow cellularity may underestimate the functional deficit. Second, the erythroid liability of otherwise distinct agents scales with the replication stress they impose on maturing erythroblasts, a variable that could in principle be managed by scheduling or by protecting the terminal maturation compartment rather than by changing target class. That intermittent dosing of camonsertib mitigates anemia without compromising antitumor efficacy (*14*) is consistent with a maturation compartment that recovers when the perturbation is withdrawn.

Cytostatic genotoxic chemotherapy imposes a replication-stress-driven program of accelerated but abortive erythropoiesis, uncoupling erythroid differentiation from the completion of terminal maturation through a regulatory state in which the amplifying divisions that build erythroid identity are curtailed. Recognizing chemotherapy-associated anemia in part as a replication-stress-driven maturation defect suggests new ways to anticipate, monitor, and potentially mitigate this common toxicity

Our study contains some limitations: our primary system uses cord-blood CD34+ cells, which are fetal-like and may differ from adult bone marrow progenitors in their stress responses and globin output; extension to adult HSPCs and to in vivo models will be important, and we make no claim about patient bone marrow on the basis of these data. Further, our single-cell analyses describe how the trajectory is reshaped and identify candidate regulons, but establishing causality for individual transcription factors will require perturbation.

## Materials and methods

### Enrichment and isolation of cord blood-derived (CB-)CD34+ cells

Cord blood bags were obtained from University Hospital Basel under informed consent, approved by the ethics committee Nordwest-und Zentraschweiz (EKNZ), with the permission number: BASEC-Nr. 2025-00533. Enrichment and isolation of CD34+ cells was done using the EasySep™ Human CD34 Positive Selection Kit II with EasySep (STEMCELL #17856) and following manufacturer’s instructions unless specified. Briefly, cord blood bags containing 60 to 100 mL of blood were brought up to room temperature. Blood was transferred to a 125 mL flask and blood was diluted 1X in recommended medium (Ca^2+^, Mg^2+^-free DPBS (ThermoFisher # 14190144) supplemented with 2% FBS and 1 mM EDTA). 30 mL of diluted blood was poured per pluriMate II Tubes for density gradient centrifugation (pluriSelect #44-10050-05) pre-filled with PBMC Spin density gradient medium (pluriSelect # 60-00092-12), and consequent steps were followed according to the guidelines. After a final wash in recommended media, cells were resuspended in FBS, supplemented with 10% DMSO and aliquots were cryo-preserved. CD34 enrichment was assessed with flow cytometry, using CD34, CD38, and CD45 antibodies (Supplemental Fig. 1C). For flow cytometry protocol, see below.

### Erythroid expansion and maturation of CD34+ cells

CD34+ cells were thawed for 1 minute in a water bath at 37°C, and resuspended and washed in 10 mL of X-VIVO® 20 Serum-free Hematopoietic Cell Medium (Lonza # 04-448Q). Cells were resuspended in 1 mL of StemSpan™ SFEM II (STEMCELL # 09605). Differentiation medium (DiffF) consisted of SFEM II medium supplemented with 1% P/S. StemSpan™ Erythroid Expansion Supplement (STEMCELL #02692) was added, at 50% of the manufacturer’s recommended working concentration, diluting it 200 times (0.5X) instead of 100 times (1X). Differentiation protocol was followed according to the manufacturer’s instructions. Cultures were maintained at 37°C, 5% CO₂ throughout expansion and maturation. Expansion phase: 10,000 to 100,000 CD34+ cells were seeded per milliliter of DiffM on day 1. On day 3 (48 hours post seeding), an equal volume of DiffM was added. On day 7, 10, and 13, cells were either harvested and replated in fresh DiffM volume at a density of 100,000 cells per mL or collected for analysis. On day 13 or 14, cells were harvested and collected for analysis, or plated in maturation media (MatM), which consisted in SFEM II media supplemented with 3U/mL EPO (STEMCELL #78007.1), 3% Human AB serum (Merck # H4522-20ML) and 1% P/S at a density of 500,000 cells per mL. On day 16, 19, and 21, cells were collected for analysis.

### Drug treatments

All the drugs were purchased from MedChemExpress. Topotecan hydrochloride (#HY-13768A), Doxorubicin hydrochloride (#HY-15142), Ceralasertib (#HY-19323), AZD1390 (#HY-109566) and Olaparib (#HY-10162), Nedisertib (#HY-101570). With the exception of Hydroxyurea (#HY-B0313-500mg), all agents were purchased as 10 mM stocks resuspended in DMSO. A 650 mM Hydroxyurea solution was prepared by dissolving 500 mg of HU in 10.2 mL of sterile water. Main stocks were kept at -20°C. Drug stock solutions were prepared fresh by diluting in IMDM to prepare 100X working stocks. These were added to the culture medium at 1% (v/v) of the final culture volume to achieve the desired final drug concentrations. Concentrations used for maturation and scRNA-seq for Topotecan, Olaparib and Ceralasertib were 20 nM, 2.5 uM and 620 nM, respectively. Drug concentrations should be titrated at the intended seeding density, as drug response was density-dependent in our hands. DMSO should never go above 0.1% w/v.

### Flow Cytometry analysis of CD34+ HSPC-derived cultures (refer to Supplementary table 1)

25,000 to 100,000 cells were collected in v-bottom plates (Corning/Sarstedt) using 400 x g for 3-5 minutes at room temperature in a benchtop centrifuge. Cells were washed with 1X PBS twice, then stained with a viability dye, depending on the antibody panel, for 10-20 minutes at room temperature. Following viability dye staining, cells were washed once with PBS, and once with FCS, followed by staining in an antibody cocktail (see concentrations below) prepared in FCS. Hoechst staining was done at room temperature, adding a drop in 2 mL of FCS containing the antibodies for extracellular staining. Extracellular staining was done for 30 minutes at room temperature. Cells were then washed with FCS twice, before analyzing them with a BD LSRFortessa. FCS files were analyzed using FlowJo version 10.

### Single-cell RNA-seq

Single-cell RNA-seq libraries were generated using the GEM-X Flex v2 for Multiple Samples kit according to the manufacturer’s protocol for cell fixation, storage, probe hybridization, sample pooling, GEM generation, and library preparation, unless otherwise specified. Briefly, 100,000 cells per sample were fixed overnight (14-18 hours) at 4°C, following the 10X demonstrated protocol "Fixation of Cells & Nuclei for GEM-X Flex Gene Expression" (#CG000782). Cells were processed in a 96-well V-bottom plate, and volumes were adjusted accordingly (to never have more than 250 μL). After quenching, cells were stored in 200 μL chilled Quenching buffer B, 20 μL pre-warmed enhancer, and 50 μL 50% glycerol. Fixed cells were stored in PCR tube strips at -80°C. Thawed fixed cells were processed using the 10X user guide "GEM-X Flex v2 for Multiplexed samples" (#CG000834, Rev C). Prior to probe hybridization, samples were normalized by cell number and pooled for subsequent GEM generation and barcoding. 16-20 samples were pooled after probe hybridization into each GEM. After PCR, library size was assessed with an Agilent TapeStation 4150 machine, using TapeStation D5000 high sensitivity reagents (Agilent #5067-5585). Prior to sequencing, an end-polishing step was added, to process 50 ng of input DNA (Flex library) using high-quality P5/P7 primers, together with the Equinox amplification module from Watchmaker Genomics applying 5 PCR cycles. Libraries were sequenced using AVITI High 75 bp (2x75) Freestyle kit at the Genomics Facility, ETH Zurich in Basel.

### Bioinformatics analysis of single cell RNA-seq

Fastq files were processed using the 10X Genomics CellRanger multi pipeline (version 10) in a Unix environment, aligning to the GRCh38-2024-A reference genome. Libraries were generated using 10x Genomics Flex chemistry, which requires the use of the multi pipeline for demultiplexing. Sequencing depth ranged from 8,000–20,000 reads per cell across samples. Subsequent analysis was performed in an RStudio environment (version 2026.05.0) with R version 4.5.3. Count matrices were processed and analyzed with Seurat (version 5.5.1 (*32*)) using default parameters, unless specified otherwise. Cells were filtered using Seurat’s default quality control thresholds; no explicit doublet detection or ambient RNA correction was performed.

Data were normalized using the LogNormalize method, and variable features were identified using Seurat’s default parameters (FindVariableFeatures, vst selection method). Dimensionality reduction was performed using principal component analysis (PCA), and the first 20 principal components were used for downstream clustering (FindNeighbors) and clustering was performed using the Louvain algorithm (FindClusters). Cell types were annotated by reference mapping to BoneMarrowMap (*19*). The presto package (version 1.0.0) was used to run the FindAllMarkers function. qs2 (version 0.2.2) was used to store and load files. SeuratExtend (version 1.2.10) and scCustomize (version 3.3.0), in addition to Seurat’s native functions, were used for plotting. Pseudotime analysis was performed using the runSlingshot function of SeuratExtend (*33*) on the PCA embedding. Gene regulatory networks (GRNs) were inferred using a customized Snakemake pySCENIC pipeline (*21*, *34*). Briefly, pySCENIC (v0.12.1) run through a Snakemake pipeline [cite or link the repository]. Raw counts from the full 82,275 dataset were used and the full workflow was run 30 times on each subset (150 runs) to account for the stochasticity of tree-based regression. In each run, co-expression modules were inferred with GRNBoost2 using the list of human transcription factors provided with pySCENIC, then pruned by motif enrichment with cisTarget (pyscenic ctx, with --mask_dropouts) using the hg38 v10 clustered motif collection (mc_v10_clust). Two gene-motif ranking databases were used: 10 kb up- and downstream of the transcription start site, and 500 bp upstream to 100 bp downstream. Motifs were annotated to transcription factors with the v10nr_clust HGNC annotation table. Regulons were aggregated across runs, retaining target genes found in atleast 5 runs , which yielded 773 regulons. Transcription factor activity was scored per cell with decoupleR (*22*) using the pySCENIC-derived regulons; ΔTF activity is the difference in mean per-cell activity between drug and control, averaged over erythroid pseudotime bins. Cell-cycle phase was assigned with Seurat’s CellCycleScoring using the Tirosh S and G2/M gene sets. Reference-mapping quality (Pass/Fail) was taken from the BoneMarrowMap mapping-error metric at the package default threshold; cells below threshold retain their nearest-neighbour label but are flagged as Fail. Module scores were computed with AddModuleScore over MSigDB hallmark sets and curated erythroid gene sets and z-scored across groups. The independent validation dataset (*5*) was reprocessed through the identical mapping and QC pipeline.

Reanalysis of the Hydroxyurea/Aphidicolin dataset (*5*) was done with the raw data deposited under GEO accession number GSE172335, samples GSM8164401 and GSM8164402. Count matrices were processed in Seurat, keeping genes detected in at least 3 cells and cells with at least 200 genes. Ensembl IDs were converted to HGNC symbols with biomaRt. Cells were annotated by day (11 or 14), treatment (DMSO, HU or APH) and donor, and doublets and unassigned cells were removed. Data were log-normalized, scaled and reduced by PCA, and the first 15 principal components were used for Louvain clustering and UMAP. Cluster markers were identified with FindAllMarkers (positive markers, average log2 fold change > 1).

Data visualization plots were generated with ggplot2 (*35*), Seurat (*32*), SeuratExtend (*33*), scCustomize (*36*).

## Acknowledgements

We thank all members of the Zaugg laboratory for helpful discussions and feedback on this work. We are grateful to the Genomics Facility Basel (ETH Zurich) and to the Flow Cytometry Core Facility of the Department of Biomedicine, University of Basel, for support with sequencing and flow cytometry, respectively. We thank the Department of Obstetrics at University Hospital Basel and the cord blood donors, without whom this work would not have been possible.

## Author contributions

J.J.L. conceived and designed the research, analyzed the data, and wrote the original manuscript draft; J.J.L. and A.K. performed research; K.D.P. contributed to the design of the single-cell RNA-sequencing approach and provided experimental guidance; J.B.Z. supervised the project; and J.J.L. and J.B.Z. obtained funding and edited the manuscript; all authors reviewed and approved the manuscript.

## Data availability and materials sharing

All data needed to evaluate the conclusions in the paper are present in the paper and/or the Supplementary Materials

The sequencing data generated in this study have been deposited in the NCBI Gene Expression Omnibus (GEO) under accession number GSE347918, accessible through the following link: https://www.ncbi.nlm.nih.gov/geo/query/acc.cgi?acc=GSE347918. The data will be made publicly available upon publication.

---

**Supplemental Figure 1.**
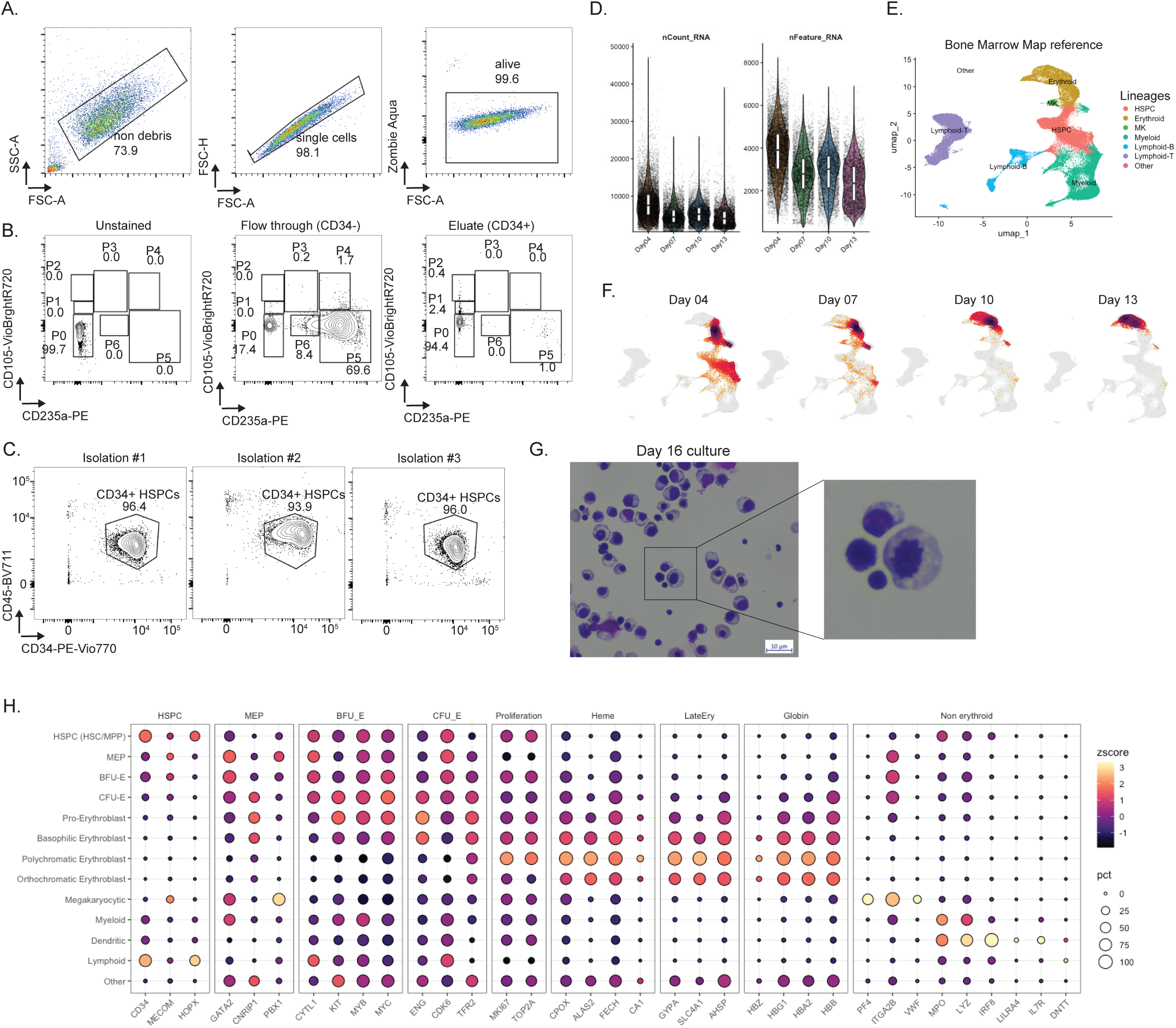
Gating strategy, CD34+ enrichment quality and transcriptional validation of the erythroid staging scheme. A. Sequential gating applied to all flow cytometry analyses in this study. Cells were gated on FSC-A versus SSC-A to exclude debris (73.9% of events), on FSC-A versus FSC-H to exclude doublets (98.1%), and on Zombie Aqua versus FSC-A to select live cells (99.6%). Numbers indicate the percentage of the parent gate. The non-debris gate is drawn permissively to retain late-stage erythroblasts, which decrease substantially in size as they mature. B. Specificity of the ENG (CD105-VioBright-R720) and GYPA (CD235a-PE) staining and position of the P0–P6 gates. Left: unstained cells, confirming that the gate boundaries capture no signal in the absence of antibody (P0 = 99.7%). Middle: the CD34-depleted flow-through fraction, containing GYPA+ cells of the input cord blood (P5 = 69.6%). Right: the CD34-enriched eluate at day 1 of culture, in which 94.4% of cells fall in P0 (ENG⌧GYPA⌧), confirming that the starting population is uncommitted. Numbers are percentages of the parent gate. C. Purity of CD34+ cells after magnetic enrichment for three independent cord-blood isolations, assessed by CD45-BV711 versus CD34-PE-Vio770. The gate marks CD34+ HSPCs; purity was 96.4%, 93.9% and 96.0%. D. Quality-control metrics for the differentiation time-course scRNA-seq dataset: total UMI counts per cell (nCount_RNA, left) and genes detected per cell (nFeature_RNA, right), by timepoint (days 4, 7, 10, 13). Boxes indicate median and interquartile range; points are individual cells.E. UMAP of the BoneMarrowMap reference atlas used for cell-type annotation, coloured by lineage (HSPC, erythroid, megakaryocytic, myeloid, lymphoid-B, lymphoid-T, other). F. Projection of the query cells onto the reference UMAP, split by timepoint (days 4, 7, 10, 13). Density shifts from the HSPC/progenitor region at day 4 towards the erythroid branch by day 13; grey points show the full reference for context. G. Dot plot of marker gene expression across BoneMarrowMap-assigned cell types, grouped by functional module: HSPC, MEP, BFU-E, CFU-E, proliferation, heme synthesis, late erythroid, globin and non-erythroid. Dot size indicates the percentage of cells expressing each gene; color indicates the z-scored mean expression across cell types. Stage-restricted expression of the markers underpinning the P0–P6 gating scheme — notably ENG, peaking at CFU-E, and GYPA, induced from the pro-erythroblast stage.

**Supplemental Figure 2.**
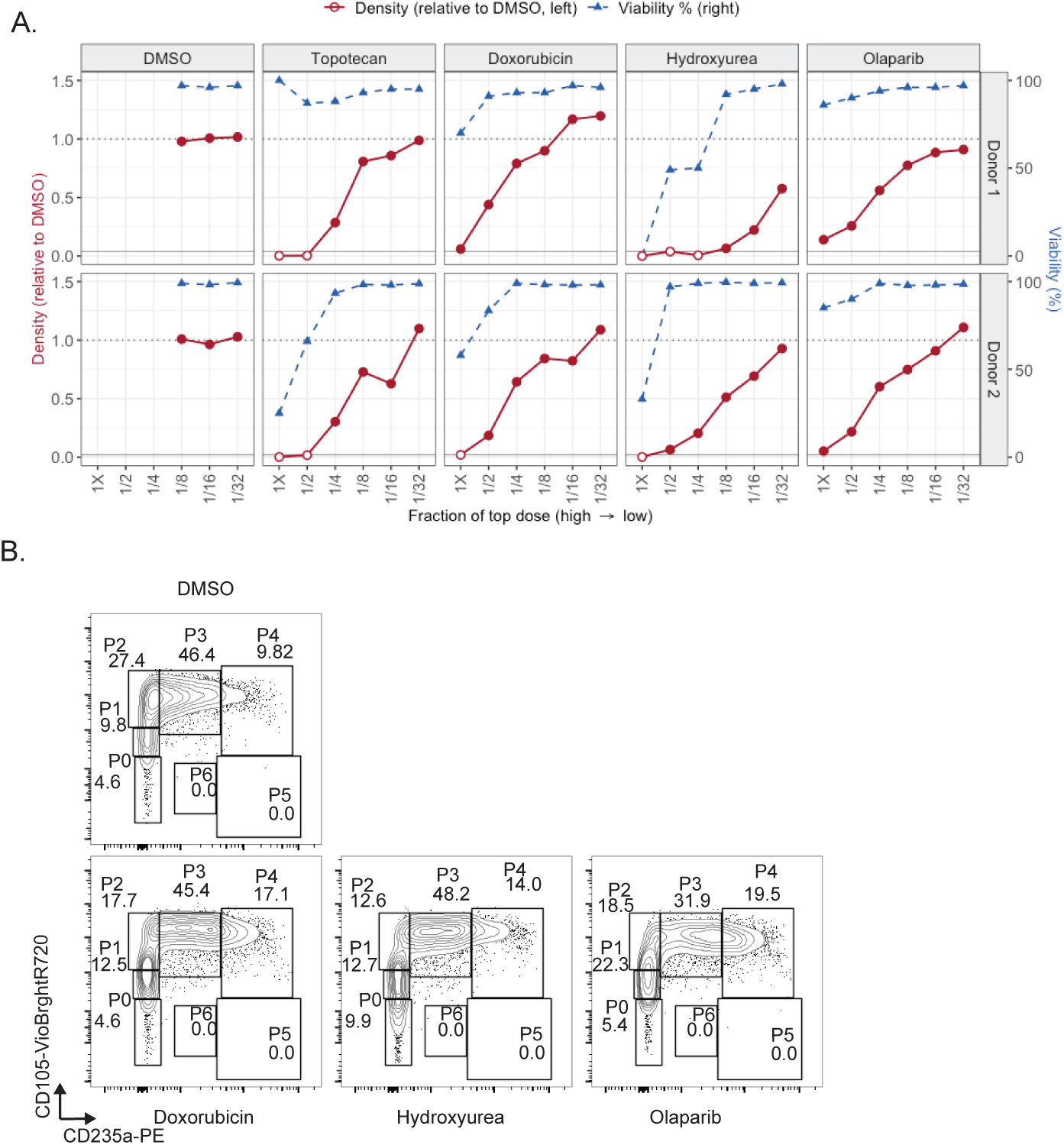
Dose–response characterisation of genotoxic agents and representative gating across compounds. A. Culture density and viability at day 7 across a six-point two-fold dilution series (1X to 1/32) for topotecan, doxorubicin, hydroxyurea and olaparib, in two independent cord-blood donors (rows). Red: density normalised within each donor to the mean of that donor’s DMSO replicate wells (left axis; dotted line = 1). Blue: viability (% live, right axis). Hollow red points indicate wells at or below the seeding density (0.04 ×10^6^/mL), where the measurement reflects failure to expand rather than a graded cytostatic effect. The DMSO panels show vehicle replicate wells rather than a dilution series, positioned along the x-axis for layout only; DMSO was added at constant volume (1% v/v) across all drug concentrations. These data define the cytostatic window used throughout the study, in which culture expansion is reduced while viability remains within 10 percentage points of control. B. Representative ENG (CD105-VioBright-R720) versus GYPA (CD235a-PE) contour plots at day 7 for DMSO (top) and doxorubicin, hydroxyurea and olaparib at the highest sub-lethal concentration (bottom), with gates P0–P6 as defined in Figure 1B. Numbers are percentages of the parent gate. Each compound reduces the P2 (mature CFU-E) fraction and increases GYPA-expressing stages (P3, P4) relative to vehicle, complementing the topotecan dose series shown in Figure 2D.

**Supplemental Figure 3.**
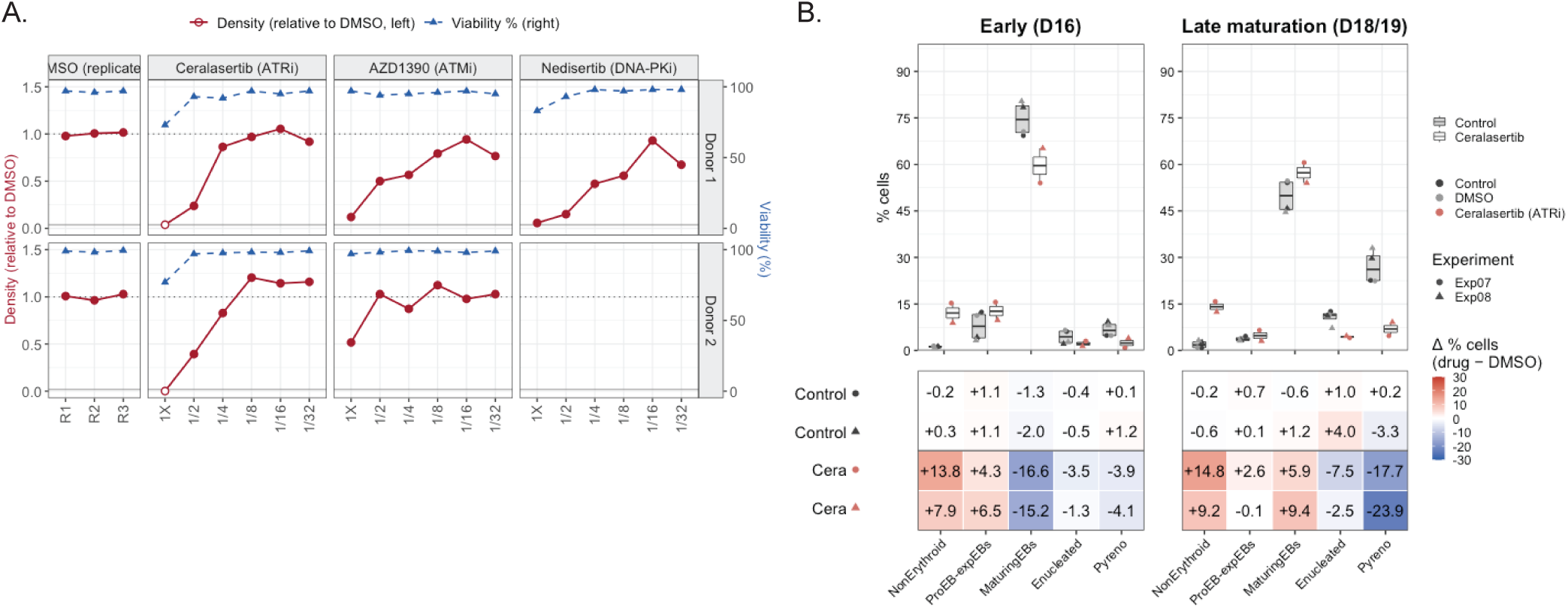
Dose–response characterisation of checkpoint kinase inhibitors and the terminal maturation defect caused by ATR inhibition. A. Culture density and viability at day 7 across a six-point two-fold dilution series (1X to 1/32) for ceralasertib (ATRi), AZD1390 (ATMi) and nedisertib (DNA-PKi). Red: density normalised within each donor to the mean of that donor’s DMSO replicate wells (left axis; dotted line = 1). Blue: viability (% live, right axis). Hollow red points mark wells at or below the seeding density (0.04 ×10^6^/mL), where the value reflects failure to expand rather than a graded cytostatic effect. The DMSO panel shows replicate vehicle wells (R1–R3) rather than a dilution series; DMSO was added at constant volume (1% v/v) across all inhibitor concentrations. Data are from two independent cord-blood donors (rows), except nedisertib, for which only donor 1 was tested (n = 1); the donor 2 panel is therefore empty. B. Maturation-phase composition under ATR inhibition, across two independent experiments (Exp07, Exp08). Gates were defined on GYPA versus DNA (Hoechst-33342) as in Figure 2E and comprise non-erythroid cells, ProEB-expEBs, maturing erythroblasts, enucleated cells (reticulocytes) and pyrenocytes; a debris gate was excluded from both the plot and the analysis. Top: percentage of cells per population at day 16 (early, left) and at the late maturation timepoint (day 18 in Exp07, day 19 in Exp08, pooled as D18/19, right), for controls (untreated and DMSO, grey boxes) and ceralasertib (white boxes). Points are individual samples; colour denotes condition and shape denotes experiment. Bottom: change in the percentage of cells per population relative to the matched DMSO control within the same experiment and day (Δ% = drug − DMSO), one row per condition per experiment; the color scale is clipped at ±30 percentage points.

**Supplemental Figure 4.**
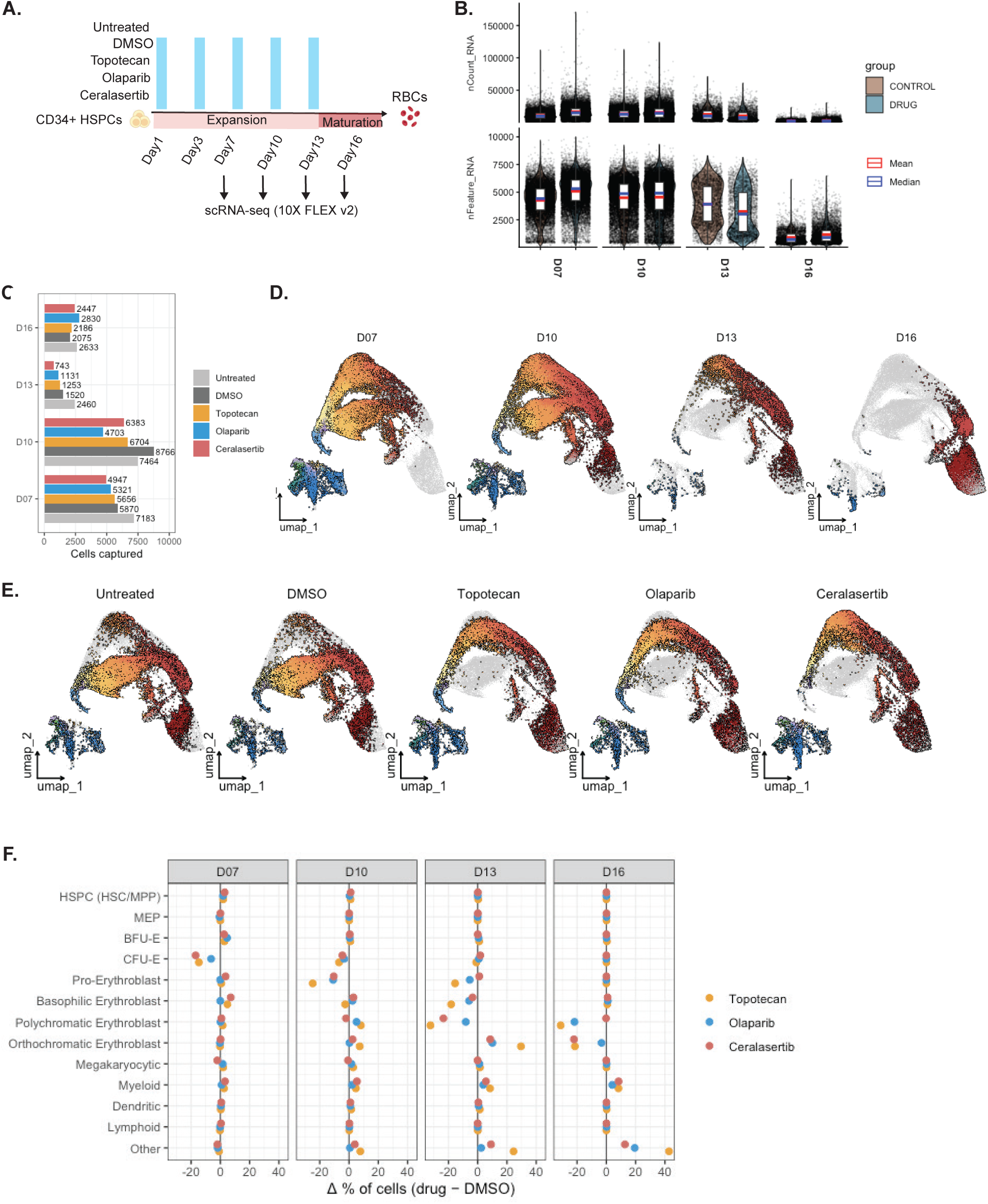
Design, quality control and composition analysis of the chemotherapy scRNA-seq time course. A. Experimental design. Cord-blood CD34+ HSPCs were differentiated under five conditions (untreated, DMSO, topotecan, olaparib, ceralasertib), with drug added from day 1 and refreshed at each medium change (blue bars). Cells were collected for scRNA-seq (10x Genomics Flex v2) at days 7, 10 and 13 during the expansion phase and at day 16, two days after the switch into maturation medium. B. Quality-control metrics per timepoint: total UMI counts per cell (nCount_RNA, top) and genes detected per cell (nFeature_RNA, bottom), split by group (CONTROL = untreated + DMSO; DRUG = topotecan + olaparib + ceralasertib). Red and blue marks indicate mean and median; points are individual cells. 82,275 cells passed quality control across all conditions and timepoints. C. UMAP of the full dataset, with cells from each timepoint highlighted in turn (colored by BoneMarrowMap-predicted cell type; remaining cells in grey). Cells progress from the progenitor-enriched region at day 7 towards the mature erythroblast branch by day 16. D. The same embedding with cells from each condition highlighted in turn. Drug-treated conditions occupy the late erythroblast region more heavily than controls at matched timepoints. E. Change in cell-type composition relative to the matched DMSO control at each timepoint (Δ% = drug − DMSO), for topotecan, olaparib and ceralasertib. Cell types are ordered along the erythroid trajectory, with non-erythroid populations and the unassigned "Other" fraction below. The three agents agree in both the direction and the approximate magnitude of change at every timepoint, despite their distinct molecular targets. Early changes are concentrated in the progenitor compartment (CFU-E depletion at day 7), while from day 13 onward the dominant change is the expansion of the "Other" fraction, which reaches approximately +43 percentage points for topotecan at day 16. Values are from a single capture per condition per timepoint, so no error estimate is available.

**Supplemental Figure 5.**
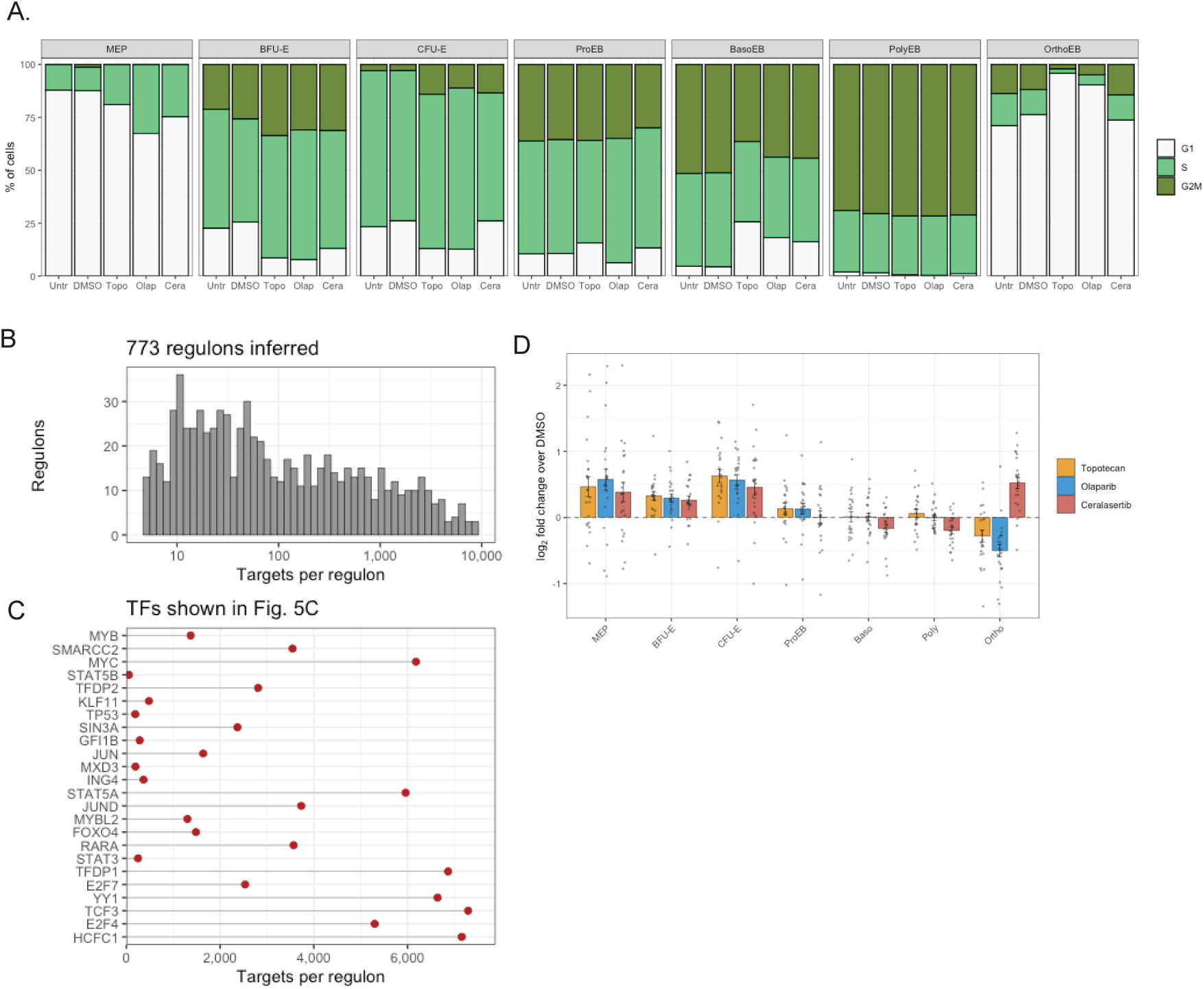
Cell-cycle composition per condition, gene regulatory network characteristics, and E2F target gene expression. A. Cell-cycle phase distribution (G1, S, G2/M) per erythroid stage for each of the five conditions, assigned with Seurat’s CellCycleScoring using the Tirosh S and G2/M gene sets. Bars are the mean percentage of cells per phase across timepoints; sample × stage combinations with fewer than 20 cells were excluded. No error bars are shown: with one capture per condition per timepoint there are no within-condition replicates, and variation across timepoints reflects differentiation kinetics rather than measurement error. The number of timepoints contributing to each bar varies by stage (MEP, 1; BFU-E, 2; CFU-E and ProEB, 3; BasoEB, PolyEB and OrthoEB, 4). Untreated and DMSO are near-identical at every stage, confirming that the vehicle has no cell-cycle effect. Relative to controls, drug-treated cells show G1 depletion with G2/M accumulation at BFU-E and CFU-E, and the reverse at BasoEB, where G1 increases. At OrthoEB, topotecan and olaparib produce a near-complete G1 accumulation (96% and 90%) with loss of both S and G2/M, whereas ceralasertib remains close to control. B. Distribution of regulon sizes across the 773 regulons inferred by pySCENIC and used for transcription factor activity scoring with decoupleR. Regulon size spans three orders of magnitude (median 60 targets, range 5 to 8,656), which should be considered when comparing activity scores between transcription factors. C. Number of target genes per regulon for the 24 transcription factors shown in Figure 5C, in the same order. D. Stage-resolved expression of canonical E2F and DREAM-complex cell-cycle target genes under each drug. Bars are the mean log2 fold change relative to matched DMSO across the gene set; error bars are SEM across genes; grey points are individual genes. The dashed line marks no change.

**Supplemental Figure 6.**
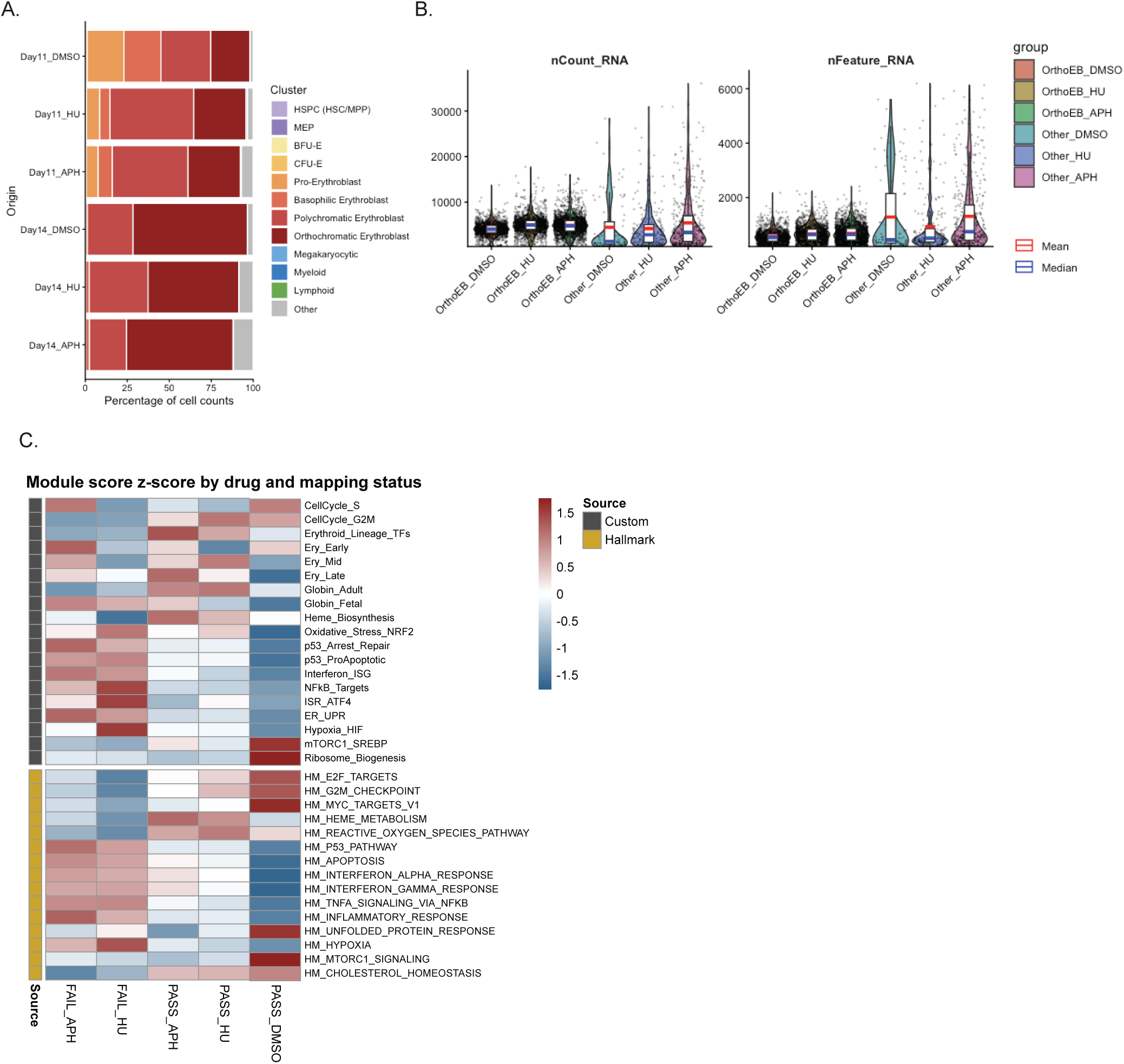
The off-reference late erythroblast state is reproduced in an independent replication-stress dataset. The PASS/FAIL classification described in Figure 6 was applied to a published dataset in which adult CD34+ HSPCs were differentiated towards the erythroid lineage under hydroxyurea (HU) or aphidicolin (APH), reprocessed through the same BoneMarrowMap reference-mapping and QC pipeline. A. Cell-type composition (percentage of cell counts, BoneMarrowMap labels) by condition and timepoint.. We note that this differentiation protocol was not carried through to terminal maturation and enucleation, so day 14 is the latest timepoint available. B. nCount_RNA (left) and nFeature_RNA (right) per group, for orthochromatic erythroblasts (OrthoEB) and unassigned ("Other") cells under each condition. Red and blue marks indicate mean and median; points are individual cells. C. Mean module-score z-scores across the six groups defined by drug and mapping status, for the same curated and MSigDB hallmark gene sets used in Figure 6E. Rows are annotated by gene-set source (custom or hallmark).

